# Spike-timing pattern operates as gamma-distribution across cell types, regions and animal species and is essential for naturally-occurring cognitive states

**DOI:** 10.1101/145813

**Authors:** Meng Li, Kun Xie, Hui Kuang, Jun Liu, Deheng Wang, Grace E. Fox, Wei Wei, Xiaojian Li, Yuhui Li, Fang Zhao, Liang Chen, Zhifeng Shi, He Cui, Ying Mao, Joe Z. Tsien

## Abstract

Spike-timing patterns - crucial for synaptic plasticity and neural computation - are often modeled as Poisson-like random processes, log-normal distribution or gamma-distribution patterns, each with different underlying assumptions that may or may not be biologically true. However, it is not entirely clear whether (and how well) these different models would or would not capture spike-timing statistical patterns across different neurons, regions, animal species and cognitive states. Here, we examine statistical patterns of spike-timing irregularity in 13 different cortical and subcortical regions from mouse, hamster, cat and monkey brains. In contrast to the widely-assumed Poisson or log-normal distribution patterns, we show that spike-timing patterns of various projection neurons-including cortical excitatory principal cells, hippocampal pyramidal cells, inhibitory striatal medium spiny neurons and dopaminergic neurons, as well as fast-spiking interneurons – all invariantly conform to the gamma-distribution model. While higher regularity in spike-timing patterns are observed in a few cases, such as mouse DA neurons and monkey motor cortical neurons, there is no clear tendency in increased firing regularity from the sensory and subcortical neurons to prefrontal or motor cortices, as previously entertained. Moreover, gamma shapes of spike-timing patterns remain robust over various natural cognitive states, such as sleep, awake periods, or during fearful episodic experiences. Interestingly, ketamine-induced general anesthesia or unconsciousness is associated with the breakdown of forebrain spike patterns from a singular gamma distribution into two distinct subtypes of gamma distributions, suggesting the importance of this spike-timing pattern in supporting natural cognitive states. These results suggest that gamma-distribution patterns of spike timing reflect not only a fundamental property conserved across different neurons, regions and animal species, but also an operation crucial for supporting natural cognitive states. Such gamma-distribution-based spike-timing patterns can also have important implications for real-time neural coding and realistic neuromorphic computing.

## Introduction

Spike-timing patterns play important roles in synaptic plasticity and neural computation (Gerstner et al., 1996; Markram et al., 1997; deCharms and Zador, 2000; Song et al., 2000; Sjostrom et al., 2001; Lisman and Spruston, 2005). During cognitions and behaviors, neurons discharge their spikes *in vivo* with tremendous variability in both the “control” resting states and across trials within the same experiments in response to identical stimuli (Shadlen and Newsome, 1994; Brown et al., 2004; Faisal et al., 2008). The irregularity of inter-spike intervals has long been suggested to be a fundamental process of cortical communication (Shadlen and Newsome, 1998; deCharms and Zador, 2000; Mazurek and Shadlen, 2002; Ma et al., 2006; Caporale and Dan, 2008; Gilson et al., 2011), and it was often modeled as a Poisson-like random process (Amarasingham et al., 2006; Beck et al., 2008), log-normal distribution (Hromadka et al., 2008; Mizuseki and Buzsaki, 2013a) or gamma distribution (Kuffler et al., 1957; Averbeck, 2009; Maimon and Assad, 2009; Mochizuki et al., 2016), each with a set of assumptions that may or may not be biologically true. A growing number of observations has shown that the neuronal spike pattern in many cortical areas seems to be inconsistent with the Poisson process, indicating that the Poisson process can either underestimate or overestimate the variabilities of neuronal spike patterns (Kara et al., 2000; DeWeese et al., 2003; Lindner, 2006; Heil et al., 2007; Kang et al., 2010; Berkes et al., 2011; Li et al., 2015; Moezzi et al., 2016).

More recently, log-normal distribution has been proposed to account for the variables of synaptic weights, the firing rates of individual neurons, the synchronous discharge of neural populations, the number of synaptic contacts between neurons and the size of dendritic boutons (Buzsaki and Mizuseki, 2014). Given the fact that statistical-distribution patterns of spike activity play important roles in determining the functions of neural circuits, it is necessary to determine what actual statistical distributions spike-timing patterns exhibit across various cell types, circuits and brain states across the evolutionary spectrum.

In the present study, we set out to investigate whether and how well log-normal distribution, gamma distribution, and/or Poisson model(s) would capture statistical features of spike-timing patterns. To maximally ensure that any variation in spike timing was not due to technical differences in terms of recording methods and/or spike-sorting qualities, we employed a large-scale *in vivo* tetrode recording in freely-behaving mice and hamsters to collect neural spike datasets, under similar experimental protocols, from 11 brain regions (10 mouse brain regions and one hamster brain region), spanning from the prefrontal cortex, primary visual cortex, somatosensory cortex and auditory cortex to the motor cortex and limbic systems - such as the hippocampus and amygdala, as well as midbrain dopaminergic neurons in the ventral tegmental area (VTA). To further expand the brain regions and animal species, we also analyzed the previously published datasets from the parietal cortex (parietal 5d) of the monkey brain and the primary visual cortex (V1) of the cat brain as a way to assess cross-species variability in spike-timing patterns.

Given the existence of distinct cell types in the brain and their possible differences in spike-timing dynamics, we separated the recorded single units into distinct cell subtypes - including excitatory cortical cells, CA1 pyramidal cells, inhibitory medium spiny neurons in the striatum, VTA dopaminergic (DA) neurons and fast-spiking interneurons - before performing statistical analyses. Finally, we investigated whether such dynamic spike-timing patterns are varied over different *natural cognitive states*, including sleep, awake-resting state and during fearful stimulation, as well as pharmacologically-manipulated *unnatural states*, such as ketamine-induced general anesthesia – an unconscious state which differs from the natural unconscious state, namely, sleep.

By systematically comparing three different models (namely, log-normal distribution, gamma distribution and Poisson) using the datasets collected from the 13 brain regions surveyed in four different animal species and under multiple different natural or unnatural brain states, we found that the spike-timing patterns of individual neurons all invariantly conformed to gamma distribution. This fundamental property is conserved across different neurons, brain regions and animal species despite large local heterogeneity. This finding will be valuable to the better examinations of real-time neural code and modeling of neuromorphic computing.

## Results

### Statistical properties of neural spike-timing patterns

Spike-timing patterns, as measured by inter-spike-interval (ISI distribution probability plot), are known to be highly variable across all stages of cognitive states, including the quiet-awake resting state (for examples, see Figure 1A, Upper Subpanel). It is well known that ISI patterns did not follow random Gaussian distribution, but exhibited a skewed distribution with heavy tails (Figure 1A, Lower Subpanel). Given that Poisson model is known to underestimate or overestimate the variabilities of neuronal spike patterns and belongs to a special case of the gamma-distribution (when shape parameter *k* equals to 1), we preceded the assessment of log-normal distribution and gamma distribution in the term of their effectiveness in capturing the spike-timing dynamics across all spike datasets. Accordingly, we conducted a two-step analysis on individual neurons’ activity patterns by asking two basic questions:

**Figure 1.**
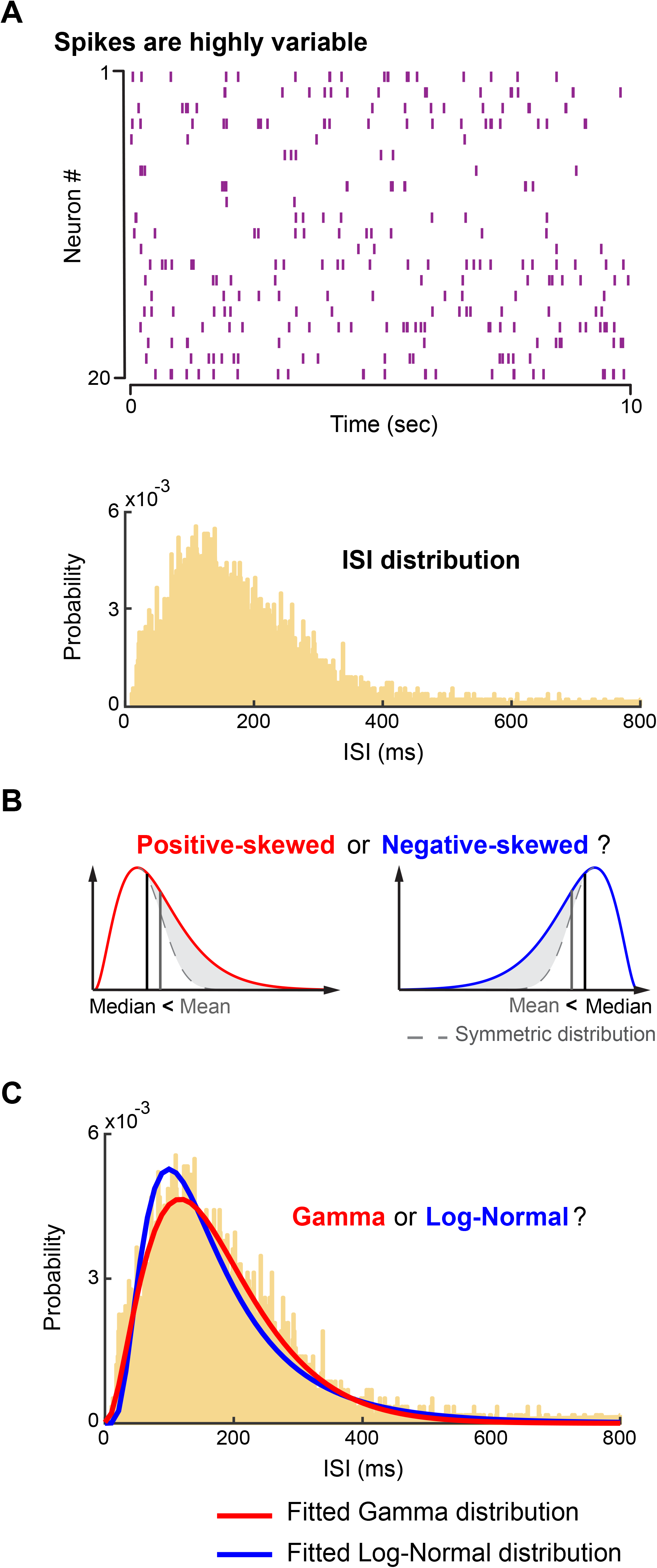
Statistical properties of neural spike patterns. **(A)** Neuronal spikes are highly variable. The upper subpanel illustrates the spike-activity patterns of 20 neurons within a 10-sec time period recorded from the VTA region of a mouse during the awake state. The lower subpanel shows the ISI distribution of a sample VTA DA neuron. **(B)** Illustrations of a positive- and negative-skewed distribution. The red and blue curves illustrate a positive-skewed distribution and a negative-skewed distribution, respectively. Gray dashed lines denote symmetric distributions. Skewness (*γ*) and nonparametric-skew (*S*) are applied to examine the positive/negative-skewed properties. (**C**) Illustrations of a gamma-distribution model and a log-normal distribution model. Fitted gamma distribution and log-normal distribution of the ISI in the lower subpanel of (**A**) are plotted as red and blue curves. The goodness-of-fit analysis is conducted to compare the gamma-distribution model with the log-normal distribution model for characterizing neuronal spike patterns.

**Step 1**: Do the neural spike-timing patterns follow a positive-skewed or negative-skewed distribution (Figure 1B)? We characterized the distribution of inter-spike intervals (ISIs) for neural spike patterns by using two well-defined statistics - namely, nonparametric-skew (*S*) and skewness (*γ*) (see Materials and Methods). In statistics, these two parameters are measurements of the skewness (or long-tailedness) of a random variable’s distribution – that is, the distribution’s tendency to lean to one side or the other of the mean (Figure 1B). A positive-skewed distribution (red curve in Figure 1B) has *S* > 0 and *γ* > 0, a negative-skewed distribution (blue curve in Figure 1B) has *S* < 0 and *γ* < 0, while a symmetric distribution (gray dotted curves in Figure 1B) has *S* = 0 and *γ* = 0.

**Step 2**: Which statistical distribution model is the best to fit the (positive) skewed distribution? As two widely-used distributions for modeling and analyzing non-negative skewed data - namely, the gamma-distribution model and the log-normal distribution model - these models were considered as alternatives for comparison (see Materials and Methods). The gamma distribution and log-normal distribution have been previously applied for modeling firing patterns of neuron population (Mizuseki and Buzsaki, 2013b) and spike patterns of single neurons (Maimon and Assad, 2009; Pipa et al., 2013).

In Figure 1C, the ISI distribution shown in Figure 1A was analyzed here as an illustrative example. It is evident that the gamma-distribution model provided more accurate-fitting results than the log-normal distribution model. It should be noted that, the parameters of both gamma and log-normal distributions in all datasets were directly estimated from the raw ISIs of neuronal spike activities by using the maximum likelihood estimates (MLE) method, which guaranteed that the analyses of ISI distributions were independent of the bin sizes used for the illustrations of ISI histograms.

We then conducted goodness-of-fit analyses to quantitatively discriminate between the gamma-distribution model and log-normal distribution model. The ratio of maximized likelihoods (RML), which is a statistical test used for comparing the goodness-of-fit of two models (Cox, 1961; 1962; 2013), was employed for discriminating the gamma-distribution model and the log-normal distribution model. The natural logarithm of RML, denoted as *D*, was calculated as the measurement of the goodness-of-fit (see Materials and Methods). That is, the gamma-distribution model precedes the log-normal distribution model if *D* > 0; otherwise, choose the log-normal distribution as the preferred model. The larger the absolute value of *D*, the more accurate fitting the result of the chosen model over the other model.

In the following sections, we conducted this two-step analyses on *in vivo* datasets of neuronal spike activity recorded from a wide range of cortical and subcortical areas under one or multiple brain states or experimental conditions in mice, hamsters, cats and monkeys.

### Principal cells exhibited gamma-distribution patterns during the quiet-awake state

We used 128-channel tetrode arrays (Xie et al., 2016) to record neurons from 10 cortical and subcortical regions - namely, anterior cingulate cortex (ACC), retrosplenial cortex (RSC), somatosensory cortex, ventral tegmental area (VTA), primary visual cortex (V1), secondary auditory cortex (2^nd^ AuV), dorsal striatum (STR), hippocampal CA1, and the basolateral amygdala (BLA) - in freely-behaving mice as well as the prelimbic cortex (PrL), a subdivision of the prefrontal cortex, of freely behaving hamsters. The qualities of recorded neurons were quantitatively measured by “Isolation Distance” (Schmitzer-Torbert et al., 2005); neurons whose “Isolation Distance” >15 were selected for the present analysis. To facilitate direct comparison, we separated putative principal cells from putative fast-spiking interneurons (see Materials and Methods). In addition, to examine the state-dependent influence on neuronal variability, we used the spike datasets collected from the quiet-awake state as animals rested in their home-cage environments. In several cases, we also collected the datasets in a set of mice when they were subjected to fearful experiences, such as earthquakes, foot shocks, and free-fall drops.

We first calculated nonparametric-skew (*S*) and skewness (*γ*) on putative principal excitatory cells, illustrated by showing datasets from the following four brain regions - namely, the ACC (neuron number n=197), the RSC (n=321), the hippocampal CA1 (n=511) and the BLA (n=451). Our results showed that all the principal cells in these regions exhibited similar positive-skewed distributions (Figures 2A-D). Skewness (*γ*) had a range of 1-10 (ACC = 3.82±0.11, RSC = 2.98±0.07, CA1 = 3.92±0.08, and BLA = 3.55±0.09, data were shown as mean ± SEM). Nonparametric-skew (*S*) were all in a range of 0.2-0.5 (ACC = 0.346±0.003, RSC = 0.335±0.001, CA1 = 0.335±0.002 and BLA = 0.358±0.003, data were shown as mean ± SEM). These results nicely demonstrated that the neuronal spike patterns in these four brain regions of mice indeed followed positive-skewed, long-tailed distributions.

**Figure 2.**
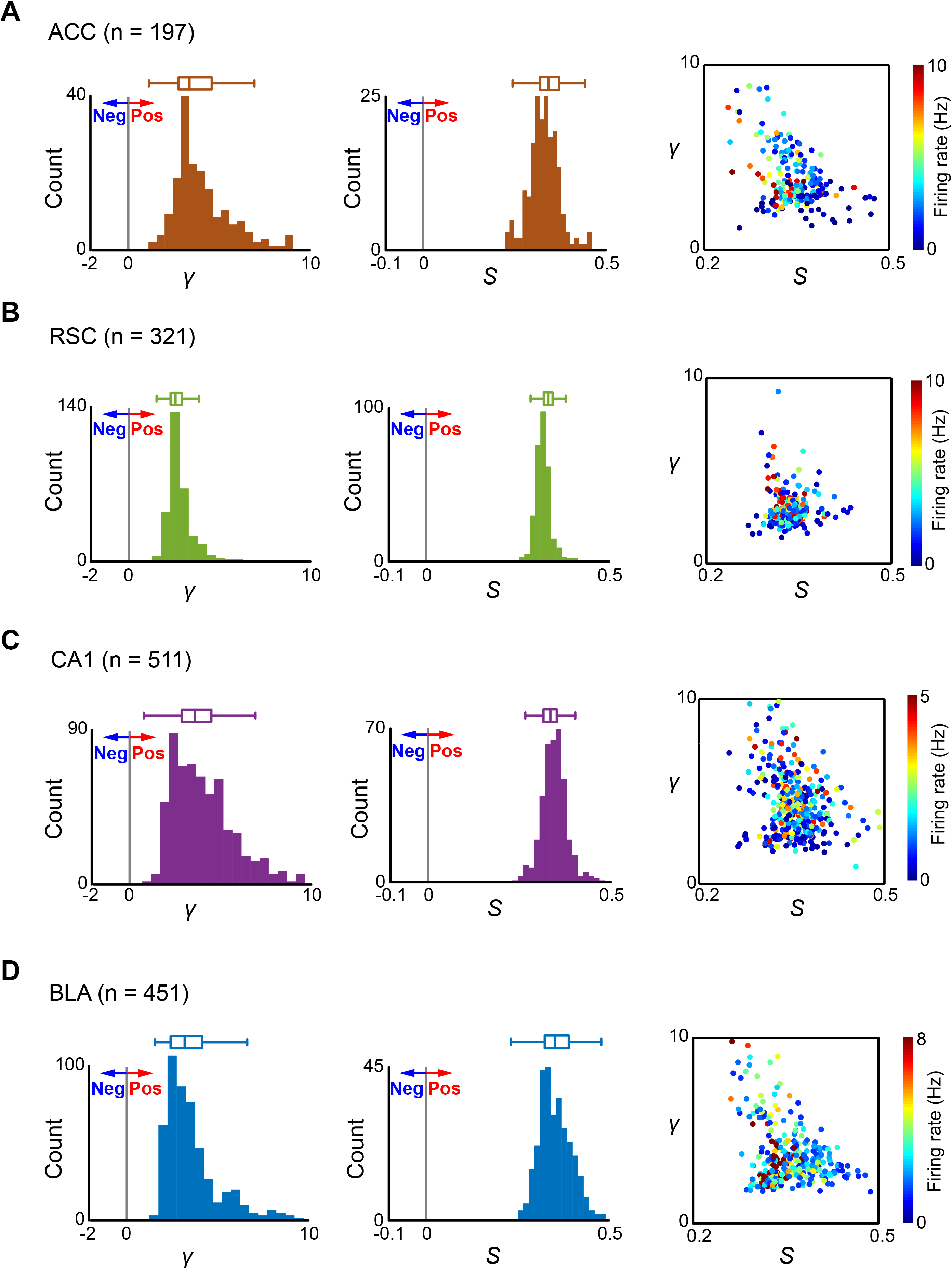
Putative principal cells in mice’s cortical and subcortical regions discharge positive-skewed, heavy-tailed spike patterns. (**A**) ACC, (**B**) RSC, (**C**) CA1, and (**D**) BLA. The left subpanels of (**A**-**D**) show the histograms of skewness *γ* in the corresponding regions, where the gray vertical lines denote the symmetric distributions (*γ*=0), positive-skewed distributions have *γ* > 0, while negative-skewed distributions have *γ* < 0. The middle subpanels of **(A-D)** show the histograms of nonparametric-skew S in the corresponding regions, the symmetric distributions have S = 0, S > 0 for positive-skewed distributions and *S* < 0 for negative-skewed distributions. The right subpanels of **(A-D)** show their relationships with the neurons’ mean firing rates, where the colors of each dot represent the mean firing rates of corresponding neurons.

We then fitted the spike patterns of each principal cell in these four brain regions during the quiet-awake state, and estimated their parameters for gamma and log-normal distributions. To avoid the under sampled error due to few data points, units with <250 ISIs were excluded in the present analysis. The left subpanels of Figures 3A-D showed fitted gamma and log-normal distributions of example principal cells for these four regions. *D* (the natural logarithm of RML) was calculated for the fitted gamma and log-normal distributions for each of the principal cells in the four brain regions. As shown in the right subpanels of Figures 3A-D, *D* > 0 was observed for all these principal excitatory cells in the four regions, demonstrating that the gamma distribution consistently outperformed on fitting neuronal spike ISIs over the log-normal distribution. We noted that the mean firing rates and *D* were highly correlated in all four regions (the correlation coefficients were: 0.80 of the ACC, 0.89 of the RSC, 0.83 of the CA1 and 0.70 of the BLA).

**Figure 3.**
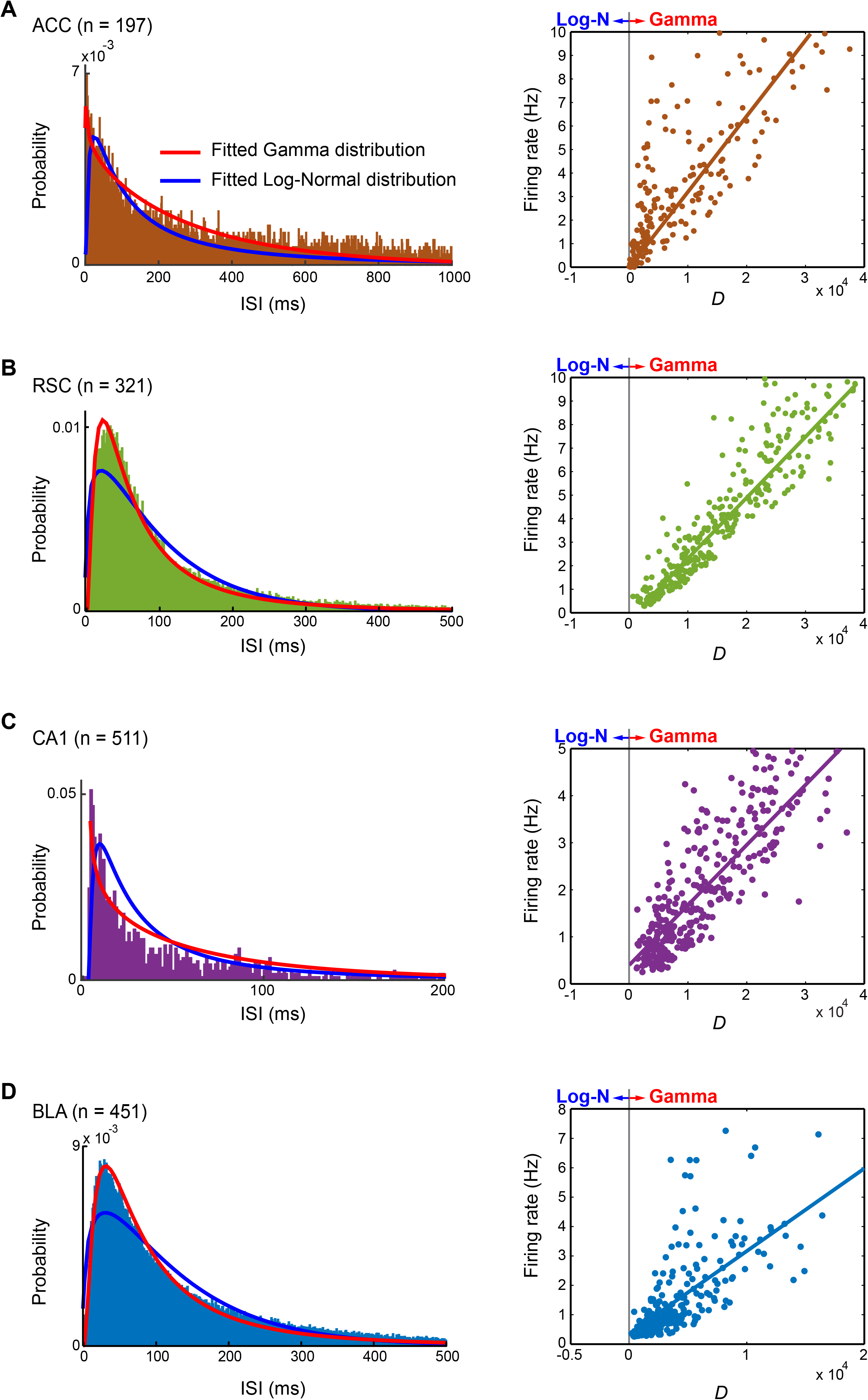
Spike-timing patterns of principal cells conformed gamma distribution across mice cortical and subcortical regions under the quiet-awake state. **(A)** ACC, **(B)** RSC, **(C)** CA1, and **(D)** BLA. The left subpanels of **(A-D)** show the spike-timing patterns of sample neurons recorded from the corresponding brain regions, their fitted gamma distributions (red curves) and log-normal distributions (blue curves). The right subpanels of **(A-D)** are the distributions of *D* (the natural logarithm of RML) vs. mean firing rates of principal cells under the quiet-awake state. The lines in these plots illustrate the linear regression between *D* and mean firing rates of principal cells.

### Principal cells exhibited gamma-distribution spike-timing patterns during sleep

Since firing patterns could vary significantly under different brain states, we asked whether the gamma-distribution model can still capture spike-timing patterns in a robust manner. As such, we compared the gamma distribution and the log-normal distribution in modeling neuronal activities of principal cells recorded from the same four mice’s brain regions during the animals’ sleep. As shown in Figure 4, the gamma-distribution model outperformed the log-normal distribution model for fitting the putative principal cells during the animals’ sleep (*D* > 0 for all four brain regions). Again, the firing rates of principal cells in these four regions were also linearly correlated with *D* under the sleep state (the correlation coefficients between firing rates and *D* were: 0.93 of the ACC, 0.71 of the RSC, 0.86 of the CA1, and 0.62 of the BLA).

**Figure 4.**
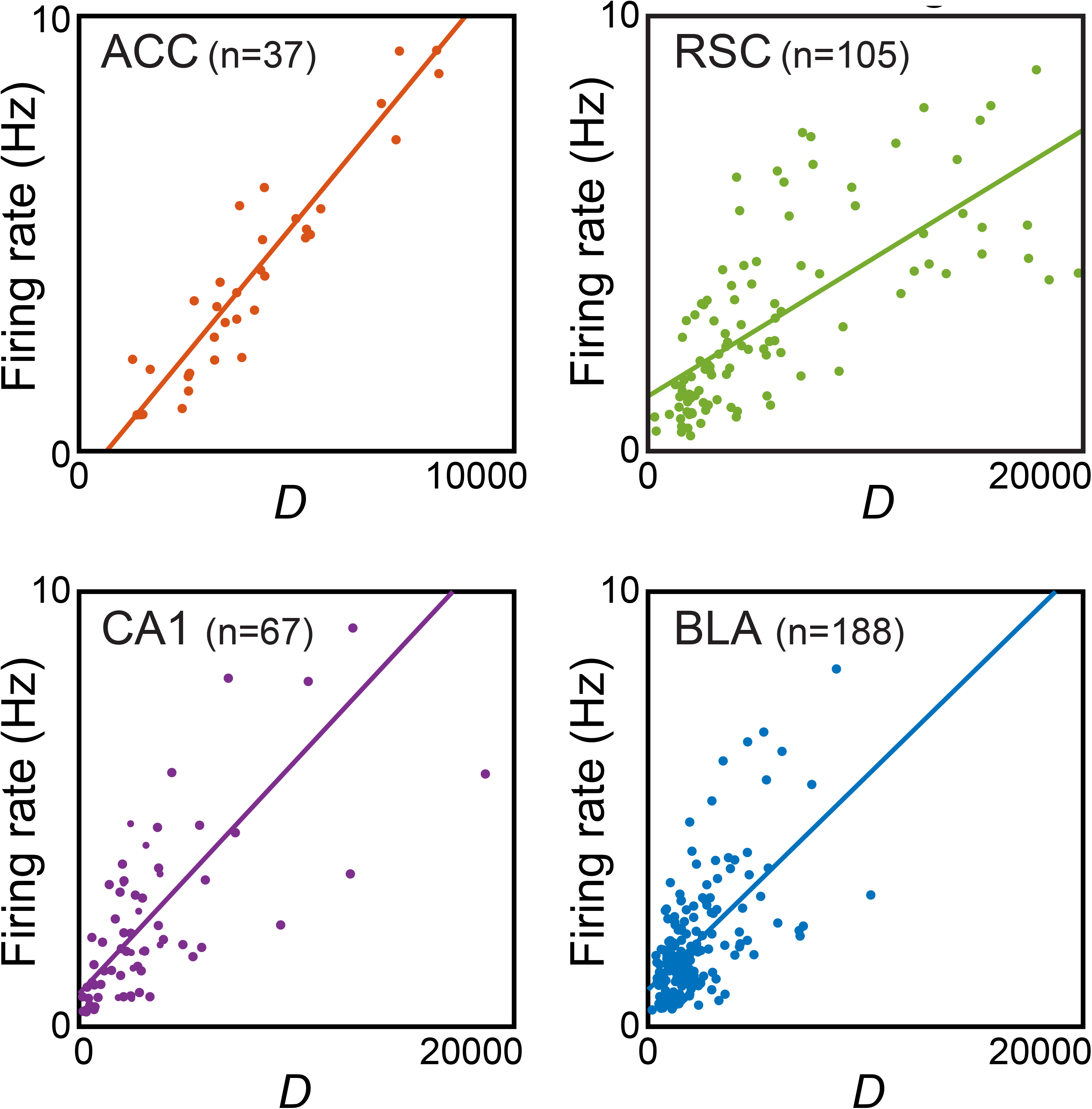
Spike-timing patterns of principal cells conformed gamma distribution across different mice cortical and subcortical regions during the animals’ sleep. The distributions of *D* vs. mean firing rates of principal cells recorded from different cortical and subcortical regions during the animals’ sleep. The lines in these plots illustrate the linear regression between *D* and mean firing rates of principal cells.

All of these analyses supported the notion that the positive-skewed distribution of neuronal spike patterns can be statistically described by the gamma-distribution model in different cortical and subcortical regions of mice under distinct brain states.

### Preserved gamma-distribution patterns across the quiet-awake and sleep states

Since the positive-skewed distribution is the intrinsic characteristic of a neuronal spike pattern, we asked whether there existed any invariant feature about this positive-skewed distribution across distinct brain states. Two parameters of the gamma-distribution model, shape parameter *k* and scale parameter *θ*, were measured under two distinct brain states (the quiet-awake vs. sleep). In the gamma-distribution model, the shape parameter *k* describes the overall envelope of an ISI distribution, while the scale parameter *θ* is related to the mean firing rate within the uncertain state (the reciprocal of the scale parameter, *r* =1/*θ*, is known as the rate parameter). The shape parameter *k* provides useful measurements for spike dynamics under different cognitive states, with a higher *k* indicating more regular spiking discharge patterns and a lower *k* indicating more irregular spike activities.

Analyses were compared the patterns of putative principal cell activities in the four brain regions under both the quiet-awake state and the animals’ sleep (34 cells from the ACC, 102 cells from the RSC, 43 cells from the CA1, and 182 cells from the BLA). The results showed that the two parameters of the gamma-distribution model were correlated between awake and sleep states in all four regions (Figure 5, correlation coefficient R between *k*_awake_ and *k*_sleep_: ACC, R=0.89; RSC, R=0.91; CA1, R=0.94; BLA, R=0.77. Correlation coefficient R between *θ*_awake_ and *θ*_sleep_: ACC, R=0.81; RSC, R=0.71; CA1, R=0.79; BLA, R=0.48. Ps <0.0001, *t*-test).

**Figure 5.**
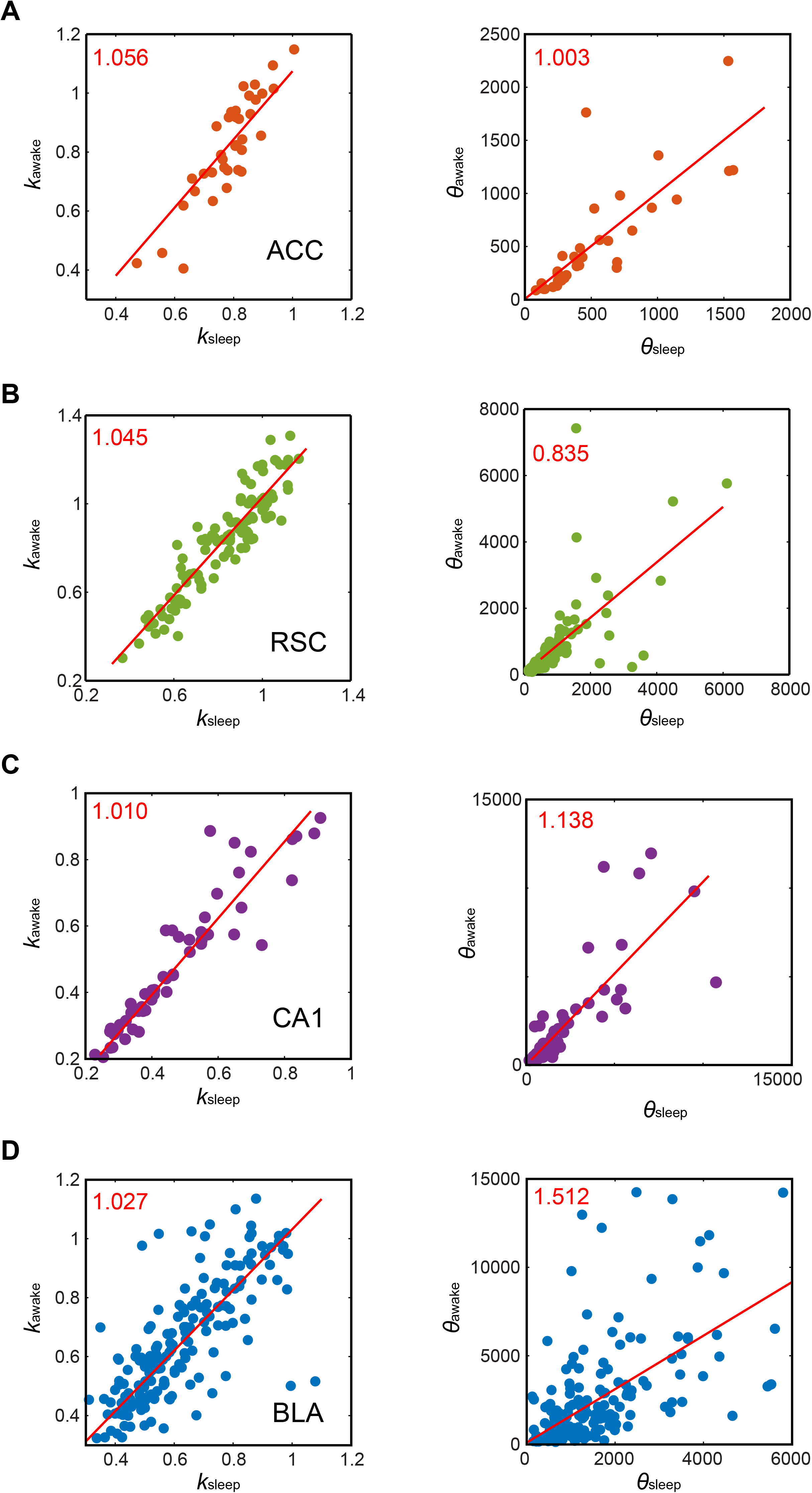
Preserved shape parameter *k* across brain states. Distributions of shape parameter *k* of principal cells in four brain regions during awake and sleep states are shown in the left subpanels of **(A-D)**, while distributions of scale parameter θ in these two states are shown in the right subpanels. Red lines denote the linear regressions of corresponding parameters, red numbers at the upper-left corner of each subpanel denote the correlation coefficients.

We further fitted these two parameters with linear functions for visually displaying the changing trends by the slope of fitted linear dependencies (shown as the red lines and red numbers in Figure 5). We observed that the transformations of the scale parameter θ (related to neuron’s mean firing rate) between awake and sleep states were varied among these brain regions. Specifically, the mean firing rates of the principal cells in the ACC remained unchanged between the two states (in the right subpanel of Figure 5A, the slopes of fitted linear dependencies was 1.003). The principal cells in the RSC exhibited increased mean firing rates in the sleep state (in the right subpanel of Figure 5B, the slopes of fitted linear dependencies was 0.835). On the contrary, as shown in the right subpanels of Figures 5C and D, the slopes of fitted linear dependences of the CA1 and BLA regions were 1.138 and 1.512, respectively, indicating that the mean firing rates of the principal cells in these two regions decreased in sleep states as compared to awake states.

Comparing the diverse transformations of *θ*, our analyses showed that the shape parameter *k* of the principal cells in all of these four brain regions was tightly correlated and highly preserved between the two states (the left subpanels of Figures 5A-D, as denoted by the slopes of fitted linear dependences of *k*: ACC, 1.056; RSC, 1.045; CA1, 1.010 and BLA, 1.027). Given the fact that the shape parameter *k* is the parameter for characterizing the overall shape of ISI distribution, our results provided strong evidence for the notion that the overall shapes of positive-skewed distributions of neuronal firing patterns were preserved across distinct brain states (sleep vs. awake), which is independent of the changes in firing rates under different cognitive states.

### Striatal medium spiny neurons exhibited gamma distribution in spike timing

Next, we investigated whether the gamma-like spike patterns reflect a general property that would remain true for GABAergic projection neurons. Thus, we employed 128-channel recording techniques and monitored activity patterns from the dorsal striatum of freely-behaving mice in the quiet-awake state. We conducted the same statistical analysis on a dataset of 294 medium spiny projection neurons (see Materials and Methods). Being the principal projection neurons of the striatum, medium spiny projection neuron is a special type of GABAergic inhibitory cells distinct from many other types of local interneurons (Preston et al., 1980; Surmeier et al., 2007).

Figure 6A showed the fitted gamma and log-normal distributions of a sample medium spiny neuron, it was evident that the gamma-distribution model produced a better result for the peak of the ISI distribution. As shown in Figure 6B, skewness *γ* = 3.77 ± 0.10 and nonparametric-skew *S* = 0.357 ± 0.003 (data were shown as mean ± SEM). These results show that the neuronal spike patterns of medium spiny projection neurons also exhibited positive-skewed, long-tailed distributions. Furthermore, the gamma-distribution model achieved superior performance on fitting neuronal spike patterns of medium spiny projection neurons over the log-normal distribution model (*D* > 0 for all medium spiny projection neurons, Figure 6C).

**Figure 6.**
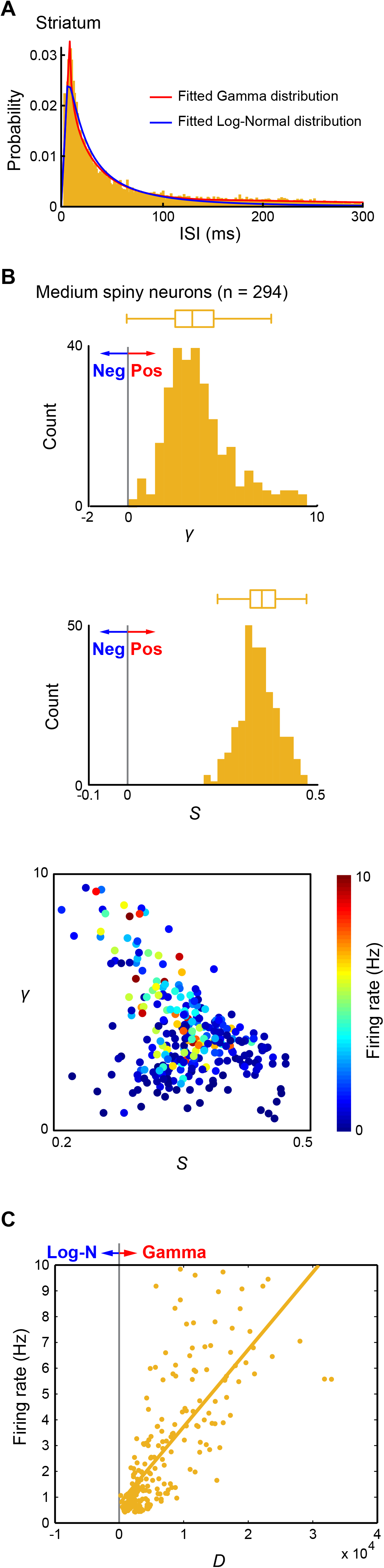
Spike-timing patterns of striatum medium spiny projection neurons conformed to gamma distribution. **(A)** The spike-timing pattern of a medium spiny projection neurons recorded from the dorsal striatum of mice, and its fitted gamma distribution (red curve) and log-normal distribution (blue curve). **(B)** Medium spiny projection neurons discharge positive-skewed, heavy-tailed spike patterns. The upper subpanel: the histogram of *γ*. The middle subpanel: the histogram of S. The lower subpanel: the distribution of *γ* and *S*, where the dots ‘colors represent neurons’ mean firing rates. **(C)** The distribution of *D* vs. mean firing rates of medium spiny projection neurons. The line in the plot denotes the linear regression between *D* and mean firing rates.

### Spike-timing patterns of fast-spiking interneurons conformed to gamma distribution

To further verify that the gamma-like spike-timing patterns were a universal characteristic of neuronal spike activity, we set out to examine if the neuronal spike patterns of fast-spiking interneurons also followed a gamma distribution. A total of 136 putative fast-spiking interneurons recorded from four mice brain regions (36 from the ACC, 31 from the BLA, 32 from the hippocampal CA1, and 37 from the RSC) were analyzed. As shown in Figures 7A and B, the neuronal spike patterns of interneurons also exhibited positive-skewed, long-tailed distributions (*γ* = 4.68±0.24, *S* = 0.327 ± 0.004, mean ± SEM). The goodness-of-fit analysis showed that the spike patterns of all these interneurons followed the gamma-distribution models (Figure 7C, *D* > 0 for all these interneurons, the correlation coefficient between *D* and mean firing rates was 0.75).

**Figure 7.**
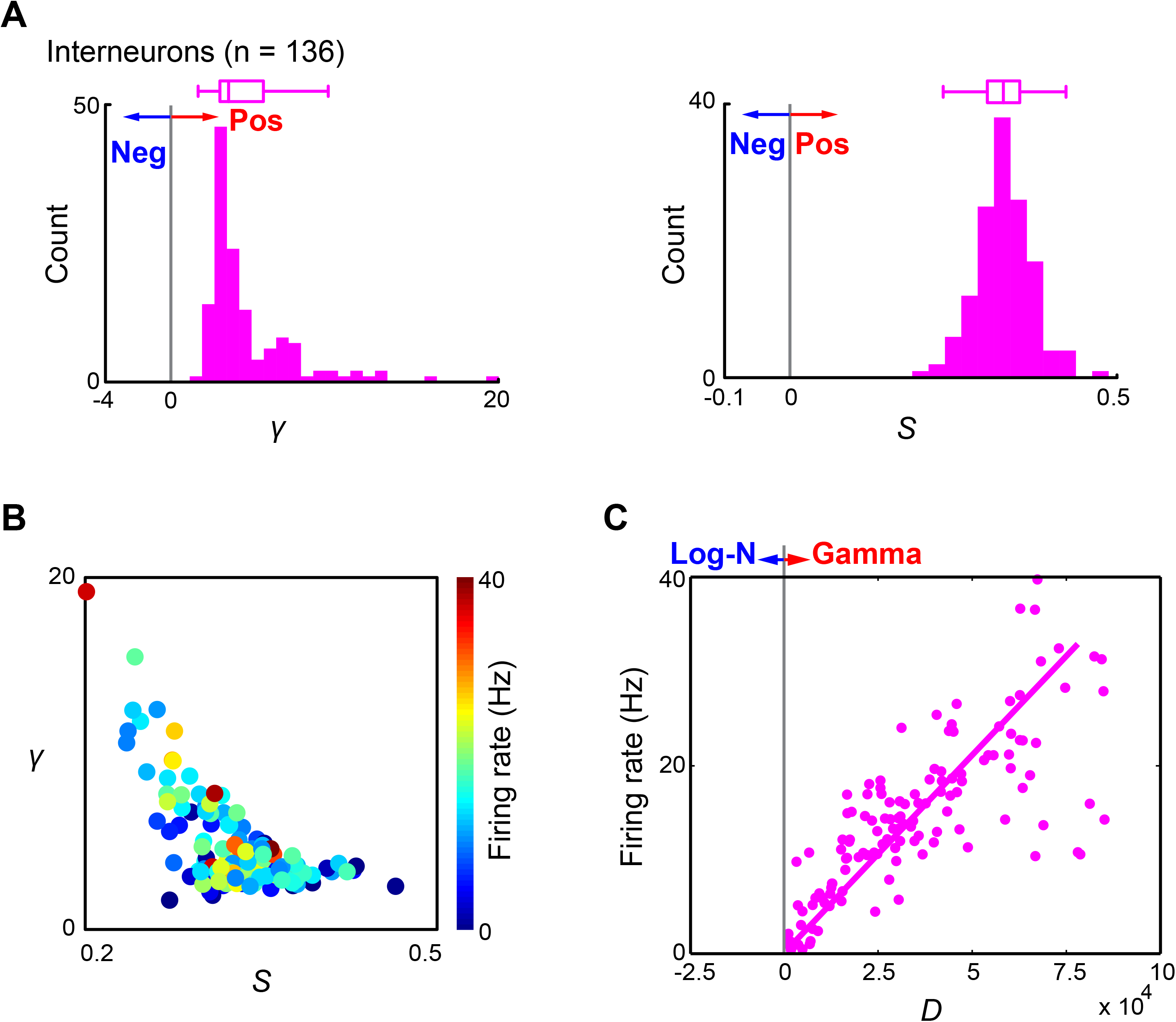
Spike-timing patterns of interneurons conformed to gamma distribution. **(A)** Interneurons recorded from four brain regions (ACC, BLA, hippocampal CA1, and RSC) discharge positive-skewed, heavy-tailed spike patterns. The left subpanel: the histogram of *γ*. The right subpanel: the histogram of *S*. **(B)** The distribution of *γ* and *S*, where the dots ‘colors represent neurons’ mean firing rates. **(C)** The distribution of *D* vs. mean firing rates. The line in the plot denotes the linear regression between *D* and mean firing rates.

### Dopaminergic neurons exhibited gamma-distribution spike-timing patterns

To further evaluate the general utility of the gamma-distribution model for different cell types, we then analyzed dopaminergic (DA) neurons from the mouse VTA region. The DA neuron dataset was collected from freely-behaving mice during the awake period using 32- or 64-channel tetrodes as we have previously described (Wang and Tsien, 2011; Li et al., 2015). DA neurons, which project over long distances to various cortical and subcortical sites (Swanson, 1982; Lammel et al., 2011; Beier et al., 2015), are well-known to subserve a wide range of biological functions, such as motivation and reward (Wise, 2004; Everitt and Robbins, 2005; Bromberg-Martin et al., 2010; Schultz, 2013), addiction (Grace, 2000; Berridge et al., 2009), fear memory and behaviors (Pezze and Feldon, 2004; Fadok et al., 2009; Wang and Tsien, 2011; Abraham et al., 2014).

We analyzed a total of 84 well-identified DA neurons, as illustrated in Figure 8A. We found that the ISI distribution of VTA DA neurons can be better characterized by the gamma distribution. Skewness (*γ* = 2.57 ± 0.21, mean ± SEM) and nonparametric-skew (*S* = 0.263 ± 0.010, mean ± SEM) demonstrated that the neuronal spike patterns of VTA DA neurons followed positive-skewed distributions with heavy tails (Figure 8B). Again, the gamma-distribution model outperformed the log-normal distribution model for fitting the neuronal spike patterns of VTA DA neurons (*D* > 0 for all VTA DA neurons, as shown in Figure 8C). And the firing rates were linearly correlated with *D* (the correlation coefficients between firing rates and *D* was 0.33).

**Figure 8.**
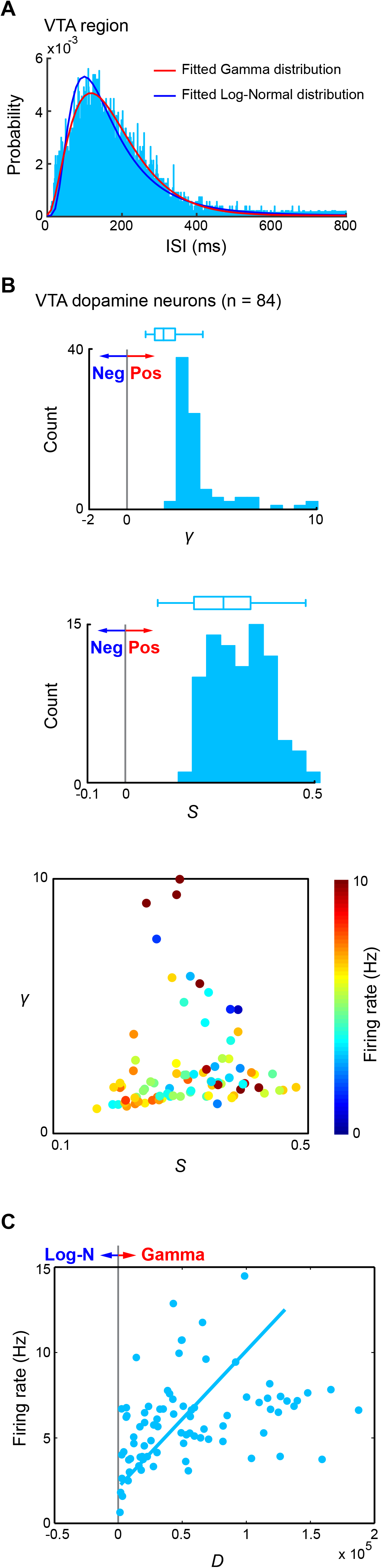
Spike-timing patterns of VTA DA neurons conformed to gamma distribution. **(A)** Fitted gamma distribution and log-normal distribution of the spike-timing pattern of a DA neuron recorded from mice VTA region. **(B)** VTA DA neurons discharge positive-skewed, heavy-tailed spike-timing patterns. The upper subpanel: the histogram of *γ*. The middle-subpanel: the histogram of *S*. The lower subpanel: the distribution of *γ* and *S*, where the dots ‘colors represent neurons’ mean firing rates. **(C)** The distribution of *D* vs. mean firing rates of VTA DA neurons. The line in the plot represents the linear regression between *D* and mean firing rates.

### Principal cells in the hamster prelimbic region also conformed to gamma distribution

To examine the generality of gamma distribution patterns across the evolutionary spectrum, we asked whether the excitatory units in the prelimbic region (PrL) of Golden Syrian hamsters would follow the gamma-distribution spike patterns. As described in our previous study (Xie et al., 2016a), we implanted 64-channel tetrode arrays bilaterally into the PrL region and recorded neural-activity patterns when animals were in their quiet-awake states. A total of 200 putative excitatory principal cells recorded from 13 hamsters were identified as stable and well-isolated units for the analyses.

The gamma and log-normal models of a sample PrL principal cell were shown in Figure 9A. The spike patterns of these cells exhibited clearly positive-skewed, heavy-tailed distributions (Figure 9B, *γ* = 3.78 ± 0.12, *S* = 0.341 ± 0.003, mean ± SEM). The goodness-of-fit analysis showed that all PrL excitatory principal cells can be better described by using the gamma-distribution models (Figure 9C, *D >* 0 for all PrL excitatory principal cells), and *D* was linearly correlated with the mean firing rates (correlation coefficient is 0.79). Together, we showed that these putative principal cells in the hamster PrL cortex also followed the gamma-distribution spiking patterns.

**Figure 9.**
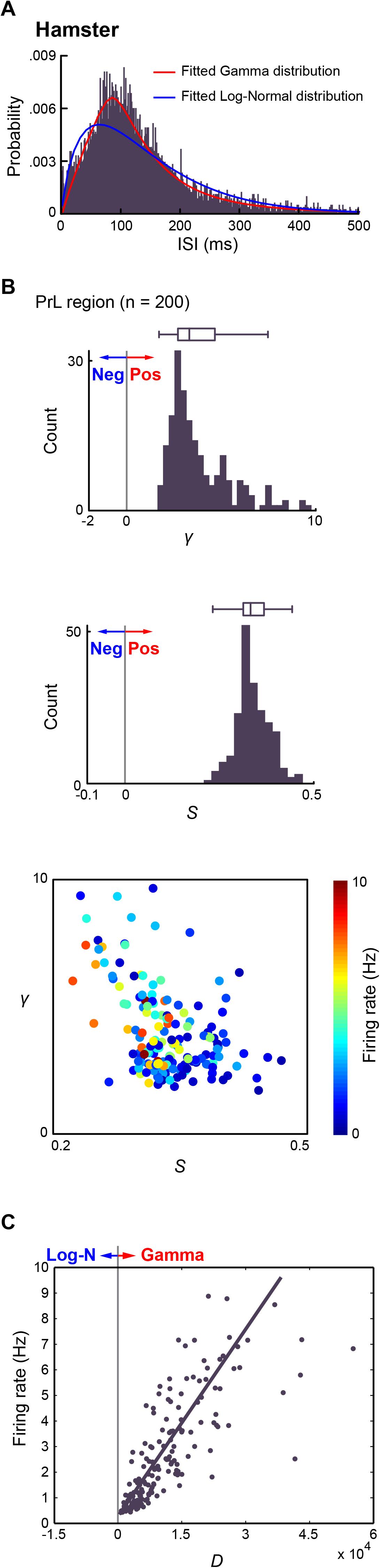
Spike-timing patterns of principal cells in hamster PrL region conformed to gamma distribution. **(A)** Fitted gamma distribution and log-normal distribution of a principal cell recorded from hamster PrL region. **(B)** Principal cells in hamster PrL region discharge positive-skewed, heavy-tailed spike patterns. The upper subpanel: the histogram of *γ*. The middle subpanel: the histogram of **(A)** *S*. The lower subpanel: the distribution of *γ* and *S*, where the dots ‘colors represent neurons’ mean firing rates. **(C)** The distribution of *D* vs. mean firing rates of principal cells in the hamster PrL region. The line in the plot represents the linear regression between *D* and mean firing rates.

### Spike-timing patterns in the monkey posterior parietal cortex area 5d exhibited gamma distribution

Next, we asked whether spike-timing patterns in primate species also followed the gamma distribution. We analyzed the datasets collected from rhesus monkeys (Li and Cui, 2013). In this study, the single-neuron activities were recorded from the parietal cortex dorsal area 5 (area 5d) on the gyral surface adjacent to the medial bank of the intraparietal sulcus, which is closely linked to movement kinematics (Hamel-Pâquet et al., 2006) and movement preparation (Cui and Andersen, 2011). Briefly, the monkeys were required to touch the fixation center at the trial beginning. Then, the first- and second- reaching goals were simultaneously displayed for 400 ms with a green square and triangle (shifted by 135° counterclockwise from the square), respectively. After a 600 ms delay, the central green dot turned off (GO signal), and the monkeys were allowed to initiate the reaching sequence to touch the locations previously cued by the square and triangle in the correct order. Throughout the trial, the monkeys were either required to maintain eye fixation (fixation condition) at the center or allowed to move their eyes freely (free-view condition). Single-reach trials were pseudo-randomly interleaved with the double-reach trials for control. Here, we analyzed a total of 211 single units and observed that the spike patterns of monkey area 5d neurons also exhibited obvious positive-skewed distributions (Figures 10A and B, *γ* =17.81 ± 1.37, *S* =0.219 ± 0.007, mean ± SEM), and can be best modeled by gamma-distribution (Figure 10C, *D* > 0 for all area 5d neurons, the correlation coefficient between *D* and mean firing rates was 0.42).

**Figure 10.**
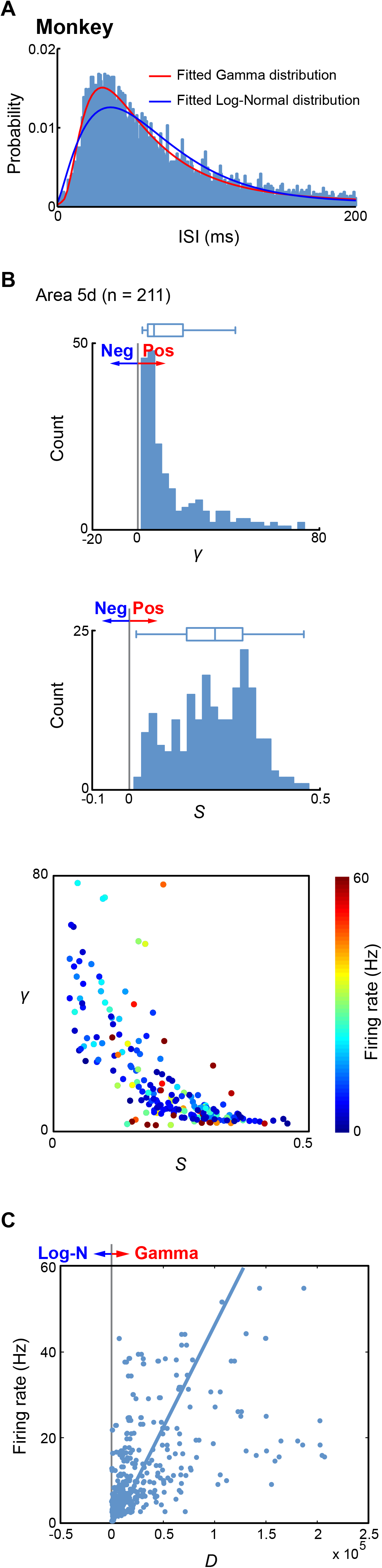
Spike-timing patterns in monkey area 5d conformed to gamma distribution. **(A)** Fitted gamma distribution and log-normal distribution of a sample neuron recorded from monkey area 5d. **(B)** Neurons in area 5d discharge positive-skewed, heavy-tailed spike patterns. The upper subpanel: the histogram of *γ*. The middle-subpanel: the histogram of *S*. The lower subpanel: the distribution of *γ* and *S*, where the dots ‘colors represent neurons’ mean firing rates. **(C)** The distribution of *D* vs. mean firing rates of neurons in area 5d. The line in the plot denotes the linear regression between *D* and mean firing rates.

### Spike-timing patterns in cat primary visual cortex conformed to gamma distribution

To further investigate the generality of gamma distribution patterns in different animal species, we analyzed a dataset of extracellular recordings from the primary visual cortex of anesthetized adult cats that were deposited in the public domain (Dan et al., 2009). This dataset was originally performed for measuring the spatiotemporal receptive fields of cortical cells (Touryan et al., 2002; Felsen et al., 2005; Touryan et al., 2005; Dan et al., 2009). One of two different visual stimuli, either 1-D white noise [473 neurons, 1-D white noise (random bars) aligned to the preferred orientation of each cell] or 2-D stimuli (202 neurons, natural images, natural phase and random phase), were delivered to the anesthetized cats. Thus, this dataset was not only ideal for verifying the gamma-distribution model, but it also afforded us an opportunity to examine if the gamma-distribution model was robust under two different types of stimulus conditions.

First, we observed that the spike patterns of the neurons in the primary visual cortex of cats all followed positive-skewed distribution under 1-D white noise (cat V1: 1-D) (Figures 11A and C, *γ* = 2.80±0.04, *S* = 0.421 ± 0.004, mean ± SEM) and 2-D stimuli (cat V1: 2-D) (Figures 11B and D, *γ* = 2.81 ±0.05, *S* = 0.392 ±0.006, mean ± SEM). Furthermore, the distributions of *γ* and *S* showed no significant differences under these two distinct types of stimuli (*p* > 0.94, *t* -test). Second, the goodness-of-fit analysis demonstrated that the gamma distribution outperformed the log-normal distribution for characterizing the spike patterns of the primary visual cortex neurons under both 1-D white noise and 2-D stimuli (Figures 11E and F, the correlation coefficients between *D* and mean firing rates were 0.44 for 1-D white noise and 0.41 for 2-D stimuli). These results demonstrated that spike-timing patterns in the primary visual cortex of cats also followed the gamma-distribution model. Moreover, overall gamma distribution remained robust over different stimulus types.

**Figure 11.**
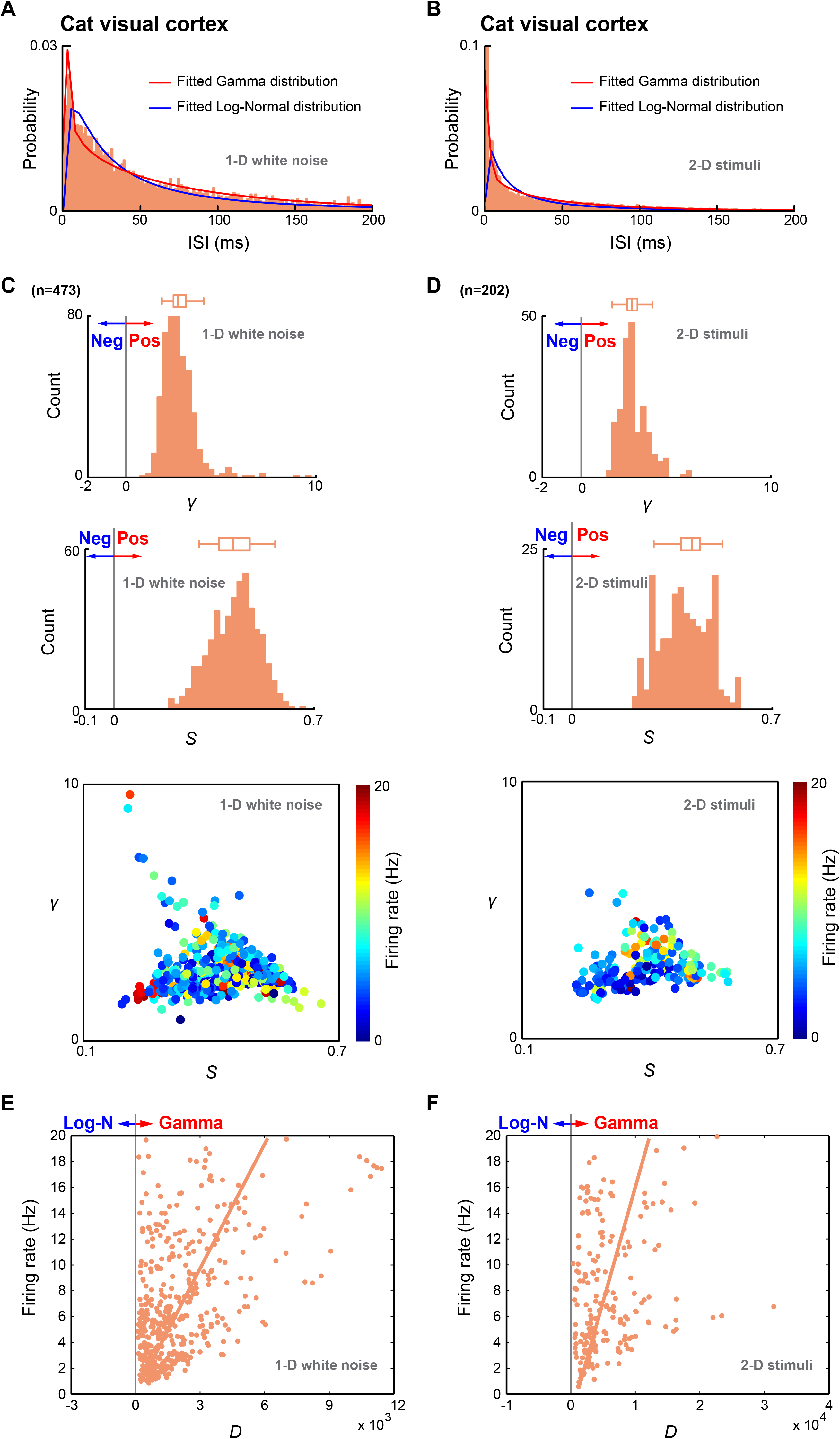
Spike-timing patterns in cat primary visual cortex conformed to gamma distribution. **(A and B)** Fitted gamma distributions and log-normal distributions of two neurons recorded from cat primary visual cortex under 1-D white noise **(A)** and 2-D stimuli **(B)**. **(C and D)** Neurons in cat primary visual cortex discharge positive-skewed, heavy-tailed spike patterns under both 1-D white noise **(C)** and 2-D stimuli **(D)**. The upper subpanels: the histograms of *γ*. The middle subpanels: the histograms of *S*. The lower subpanels: the distributions of *γ* and *S*, where the dots ‘colors represent neurons’ mean firing rates. **(E and F)** The distributions of *D* vs. mean firing rates of neurons in the cat primary visual cortex under 1-D white noise **(E)** and 2-D stimuli **(F)**. The line in the plot denotes the linear regression between *D* and mean firing rates.

### Preserved gamma-distribution spike patterns under fearful episodes

To further examine whether and how well gamma-distribution patterns capture spike-timing patterns across different brain states, we performed additional set of experiments under which mice were subjected to fearful stimulation. Specifically, two datasets were recorded from the ACC and RSC regions when mice experienced three fearful episodic events, namely, earthquakes, foot shocks and free-fall drops (see Materials and Methods) as well as during the quiet-awake and sleep states. A total of 95 and 120 principal cells were recorded from the ACC (three mice) and RSC (four mice), respectively. As shown in Figure 12, the goodness-of-fit analysis confirmed that the spike-timing patterns followed gamma distribution under these three brain states (*D* > 0).

**Figure 12.**
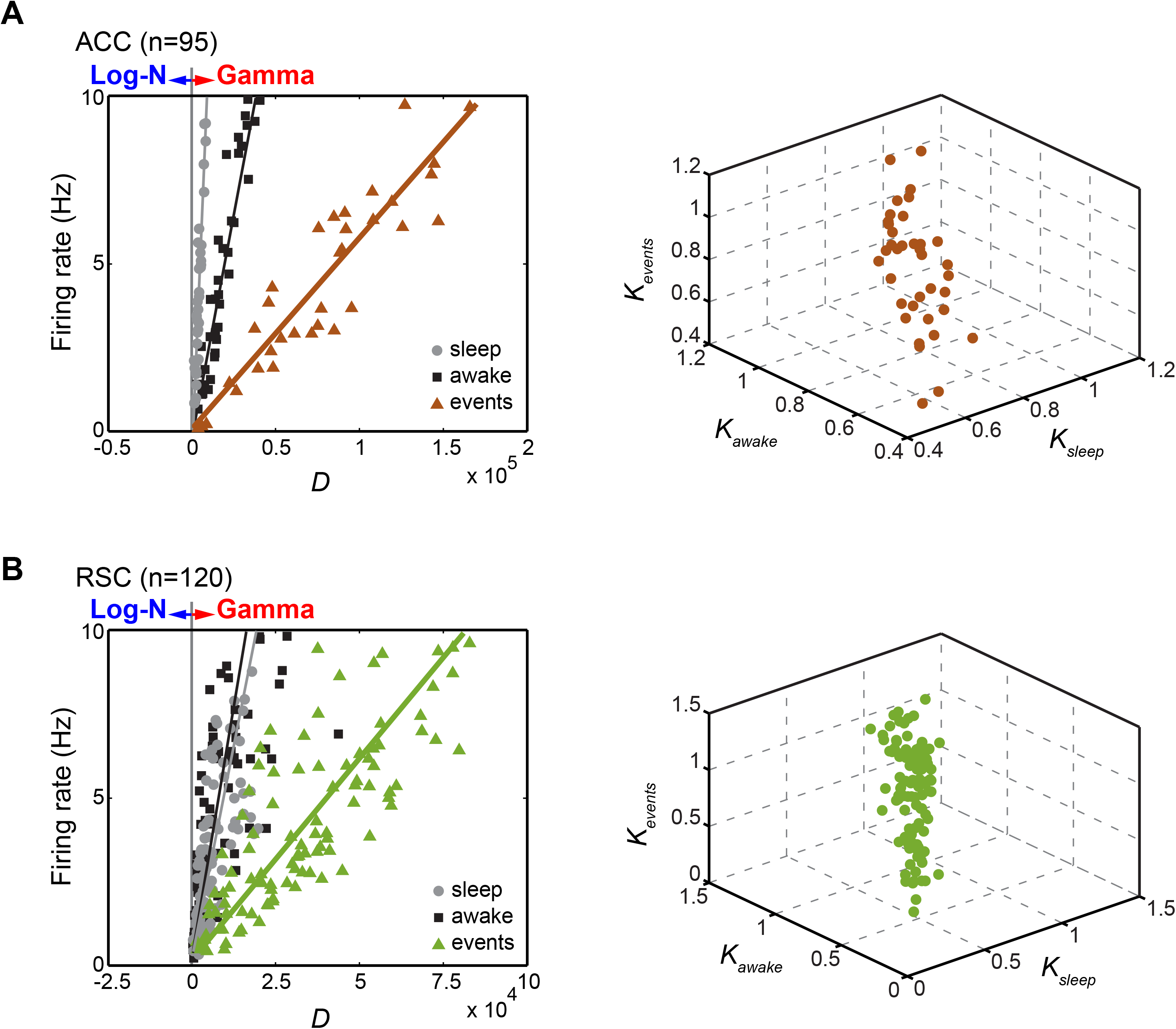
Preserved shape parameter *k* across resting states and behaving states. The distributions of *D* vs. mean firing rates of principal cells recorded from the ACC and RSC regions during two resting states (the quiet-awake state and sleep) and the behaving state are shown in the left subpanels of **(A and B)**. The lines in the left subpanels of **(A and B)** are the linear regressions between *D* and mean firing rates. 3D distributions of *k* during three states are shown in the right subpanels of **(A and B)**.

Furthermore, the firing rates and *D* were highly linearly correlated in all three brain states (left subpanels of Figures 12A and B) with *D* increasing with higher firing rates (the correlation coefficients between firing rates and *D* were: 0.94 of the ACC and 0.71 of the RSC for the quiet-awake state; 0.93 of the ACC and 0.71 of the RSC during sleep; and 0.91 of the ACC and 0.84 of the RSC during the fearful-event experiment). Interestingly, compared with the quiet-awake state and sleep, the gamma-distribution model provided even more accurate-fitting results from the stimulated states (as shown by the gradients of the fitted, linear relationships between firing rates and *D*, left subpanels of Figures 12A and B).

We also examined the robustness of the shape parameter *k* by calculating the correlations between these three distinct brain states, revealing that the shape parameter *k* was preserved not only between the quiet-awake state and sleep but also in behaving animals experiencing fearful events (right subpanels of Figures 12A and B, the correlation coefficients were: ACC, 0.83 between the quiet-awake state and sleep, 0.75 between sleep and the behaving state, and 0.71 between the quiet-awake state and behaving state; RSC, 0.96 between the quiet-awake state and sleep, 0.95 between sleep and the behaving state, and 0.94 between the quiet-awake state and behaving state). Thus, our analysis demonstrated that the spike-timing patterns conformed to gamma distribution with preserved shape parameter *k* under both resting states (the quiet-awake and sleep states) and emotionally activated states (in which fearful experiences were represented).

### Regularities of spike-timing patterns across different cells and brain regions

Given the critical roles of regularity or irregularity of spike-timing patterns in synaptic plasticity and neural computations, we further asked how spike dynamics vary over a wide range of different brain regions. Therefore, we systematically recorded and compared the regularities of neuronal spike patterns in a wide range of mouse brain regions. This includes a total of 10 mouse brain regions as follows: **1)** 84 DA neurons from the VTA; **2)** 85 excitatory principal cells from the somatosensory cortex; **3)** 615 excitatory principal cells from the RSC; **4)** 342 excitatory principal cells in the primary visual cortex (V1); **5)** 325 CA1 pyramidal cells; **6)** 41 excitatory principal cells recorded from the 2^nd^ AuV, **7)** 56 neurons from the primary motor cortex (M1); **8)** 195 excitatory principal cells from the ACC; **9)** 263 pyramidal cells in the BLA, and **10)** 221 medium spiny neurons in the STR. The datasets collected from these 10 mouse brain regions offered a unique opportunity for cross-region comparisons, because the tetrode recording techniques and criteria used for spike sorting were from the same laboratory.

Moreover, we analyzed 182 excitatory principal cells from the hamster PrL obtained in the laboratory; In addition, 475 units in the cat V1 during 1-D white noise and 207 excitatory principal cells from the cat V1 area during 2-D stimuli (from Dan laboratory), as well as 206 excitatory cells in the monkey brain area 5d (from Cui laboratory). In addition, GABAergic interneurons recorded from four mouse brain region (36 interneurons in mouse ACC region, 37 interneurons in mouse RSC region, 32 interneurons in mouse hippocampal CA1 region, and 31 interneurons in mouse BLA region) were also included in our analysis for systematically comparison of spike-timing regularities across distinct brain regions as well as different cell types. As a result, the datasets consisted of a total of 3,378 neurons obtained from 13 brain regions of the four species (mice, hamsters, cats and monkeys).

We compared the regularities of neuronal spike patterns in these brain regions by examining the shape parameters *k* of their gamma distribution patterns. Based on its definition, *k* provides an effective way to measure neuronal spike patterns’ regularities. A gamma distribution with *k*=1 is the mathematical equivalent to an exponential distribution, the corresponding neuronal spike pattern fits a Poisson process (shown as the gray horizontal line in Figure 13). Neuronal spike patterns with *k* <1 indicate that the neuron discharges more frequently in short and long intervals than the exponential distribution – namely, irregular spiking. When *k* > 1, the peaks of ISI distributions shift away from zero, and the neuron discharges more regular spiking than the Poisson process. Gamma distribution with *k* = ∞ is the distribution of no variance, and the corresponding neuronal spike pattern is perfectly regular.

**Figure 13.**
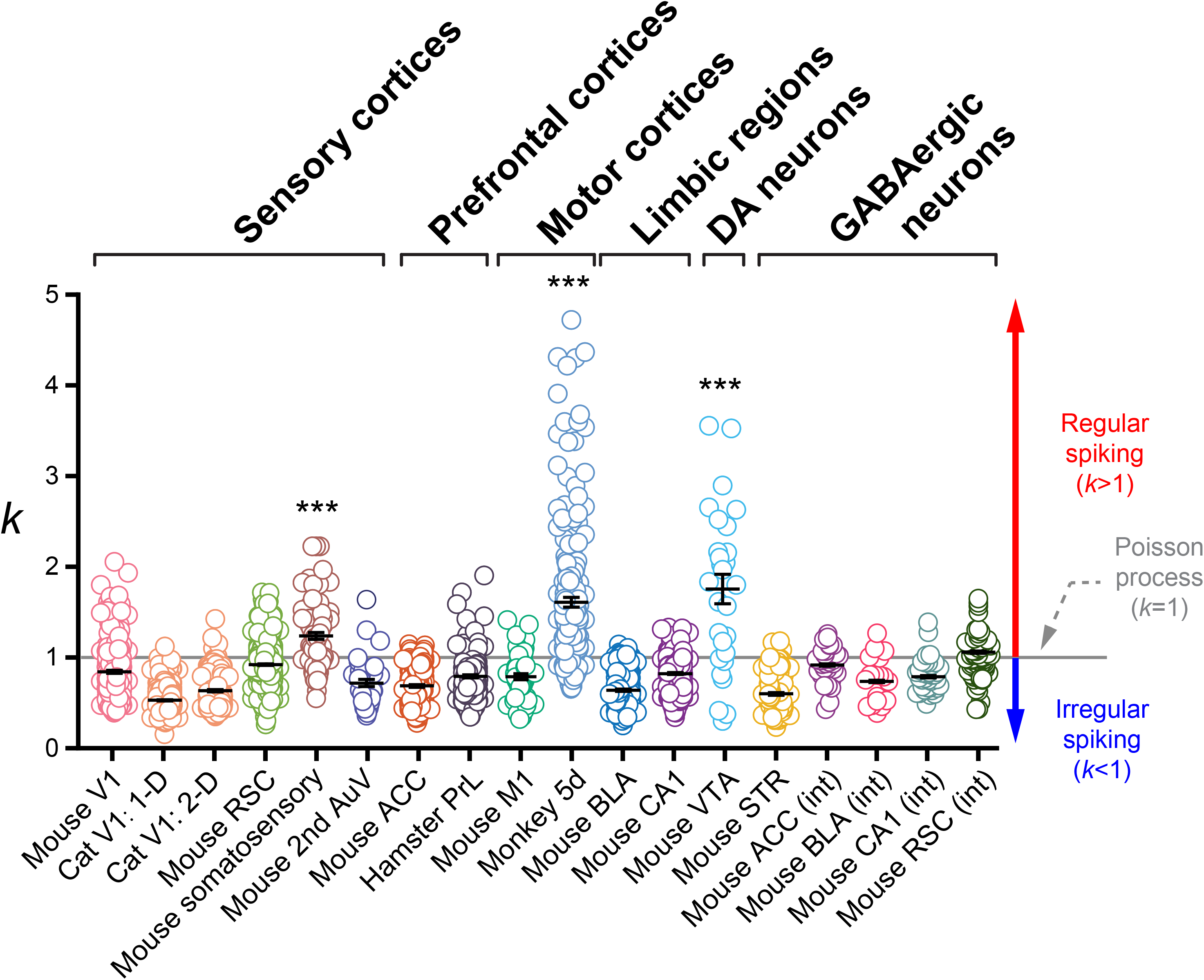
Comparisons of spike-timing regularities and irregularities across different cell types and brain regions. The distributions of *k* in 18 datasets recorded from four mammalian species (mice, hamster, cat and monkey), it consisted of a total of 3,378 neurons obtained from 13 brain regions. These neurons were grouped according to their general anatomical locations and cell types, namely, the sensory cortices (mouse V1, cat V1, mouse RSC, mouse somatosensory, and mouse 2^nd^ AuV), prefrontal cortices (mouse ACC and hamster PrL), motor cortices (mouse M1 and monkey 5d), limbic regions (mouse BLA and mouse CA1), DA neurons (mouse VTA), and GABAergic neurons (mouse STR medial spiny neurons, and fast-spiking interneurons in mouse ACC, BLA, CA1 and RSC). Each dot represents a neuron. The gray horizontal line denotes the Poisson process (k = 1), regular spiking (k > 1) and irregular spiking (*k* < 1) are shown as a red-arrow line and blue-arrow line, respectively. Error bars represent mean ± SEM.

We grouped these neurons according to their general anatomical locations and cell types, namely, seneory cortices (mouse V1, cat V1, mouse RSC, mouse somatosensory, and mouse 2^nd^ AuV), prefrontal cortices (mouse ACC and hamster PrL), motor cortices (mouse M1 and monkey 5d), limbic regions (mouse BLA and mouse CA1), DA neurons (mouse VTA), and GABAergic neurons (mouse STR, interneurons in mouse ACC, BLA, CA1 and RSC).

As shown in Figure 13, we observed that the neuronal spike patterns in most brain regions exhibited more irregular spiking than the Poisson process (*k*: 0.844±0.015 in mouse V1, 0.530±0.007 in Cat V1: 1-D, 0.636±0.013 in Cat V1: 2-D, 0.921±0.008 in mouse RSC, 0.719±0.040 in mouse 2^nd^ AuV, 0.689±0.015 in mouse ACC, 0.793±0.018 in hamster PrL, 0.791±0.033 in mouse M1, 0.632±0.013 in mouse BLA, 0.825±0.010 in mouse CA1, 0.601±0.014 in mouse STR, 0.921±0.032 of interneurons in mouse ACC, 0.749±0.061 of interneurons in mouse BLA, and 0.804±0.029 of interneurons in mouse hippocampal CA1, data were shown as mean ± SEM).

We also noted that neurons in the other three brain regions seemed to discharge more regular spikes than the Poisson process (*k*: 1.240±0.039 in the mouse somatosensory cortex, 1.610±0.055 in the monkey brain motor area 5d, and 1.806±0.104 by DA neurons from the mouse VTA, data were shown as mean ± SEM). There were significant differences between three regular-spiking regions in comparison to 15 regions which showed irregular-spiking (*p* < 0.0001, One-way ANOVA coupled with Tukey’s posthoc test). Overall, there is no obvious tendency in increased firing regularity from the sensory and subcortical neurons to prefrontal or motor cortices.

### Gamma-distribution spike patterns were dramatically altered under anesthesia

If the gamma-distribution spike-timing patterns reflect a fundamental and conserved property for natural cognitions produced by naturally occurring internal states and external inputs, we hypothesized that such gamma-distribution spike-timing patterns would be altered if naturally occurring cognitions were pharmacologically shut down (i.e., upon anesthesia). Accordingly, we used ketamine/domitor to induce general anesthesia and compared neuronal spike activities from the BLA, CA1 and RSC regions with those in the awake states from the same set of mice. We found that ketamine induced rhythmic spike discharge patterns as well as synchronized local field potential (LFP) oscillation In all these three brain regions (Figures 14A-C). Most notably, ketamine-induced general anesthesia produced profound delta band in LFP (Figures 14A-C, middle subpanels). while the frequencies of the rhythmic spike patterns were different in these brain regions -specifically, BLA > CA1> RSC, the spike patterns of pyramidal cells in these three regions during awake periods conformed to the gamma-distribution model (Figures 14A-C). However, the spike patterns of these pyramidal cells during anesthesia were broken down from a single gamma distribution (*p*<1E-5, one-sample Kolmogorov-Smirnov test) into two distinct subtypes of gamma distributions - as shown in the right subpanels of Figures 14A-C. The gamma shapes of the two subtypes of the distributions revealed that the Subtype-1 is highly irregular, reflecting bursting firing, whereas the second gamma distribution subtype showed highly regular, rhythmic spike-timing patterns in the range of delta frequency. This pharmacologically-based manipulation experiment demonstrated that the naturally-occurring and ongoing external and internal processes are essential for generating a singular gamma-distribution spike pattern for various neurons, whereas ketamine-induced general anesthesia were associated with the breakdown of this singular distribution pattern into two distinct subtypes of gamma-distribution spiking patterns.

**Figure 14.**
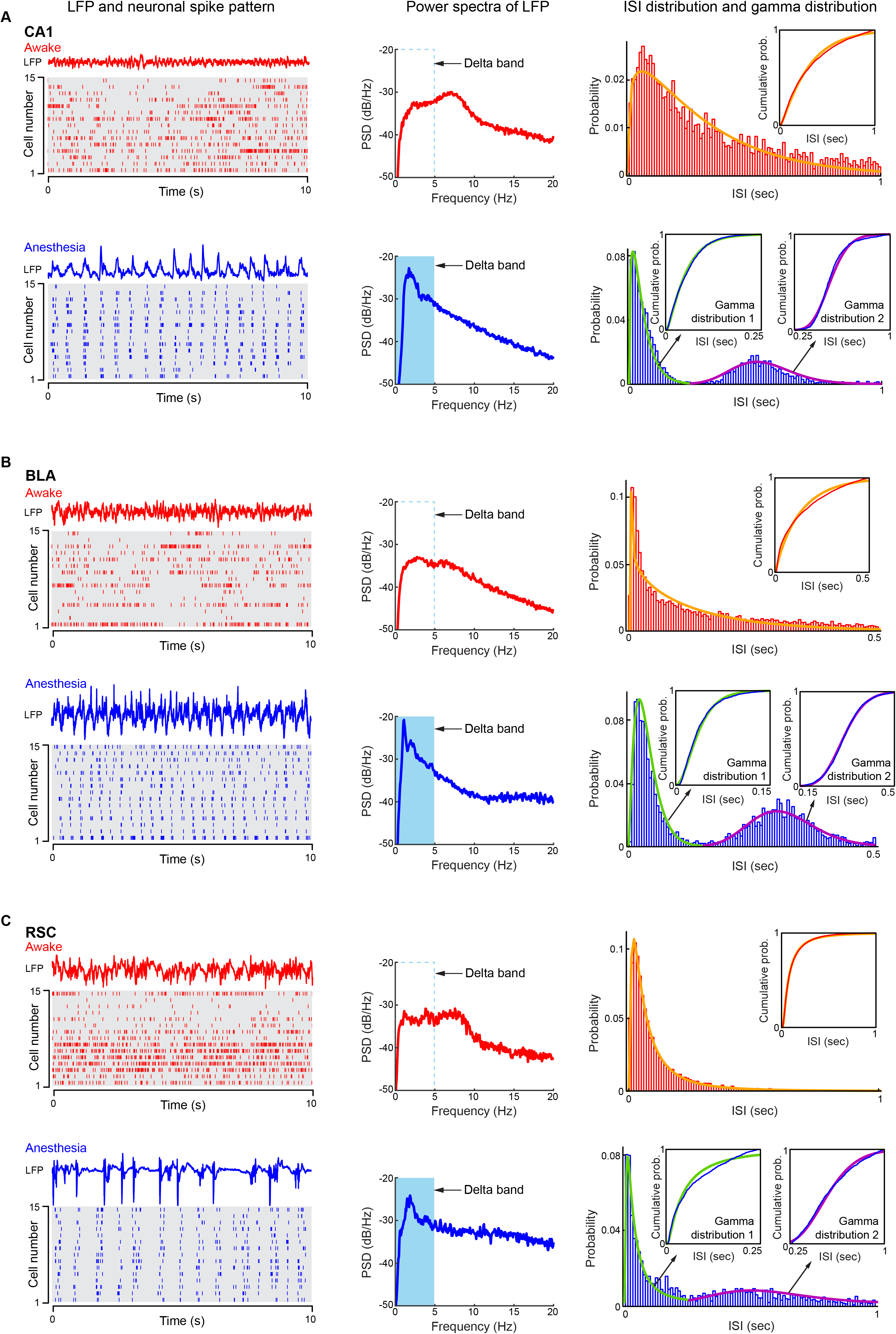
Gamma-like spike-timing patterns diminished under anesthesia. (**A-C**) Left subpanels: spike activities of 15 example principal cells recorded during awake state and anesthesia from CA1 (**A**), BLA (**B**), and RSC (**C**). Middle subpanels: power spectra of LFP recorded during awake state and anesthesia from CA1 (**A**), BLA (**B**), and RSC (**C**). Right subpanels: spike-timing patterns under the quiet awake state conform single gamma distribution (shown as the plots in red color), spike-timing patterns during ketamine-induced anesthesia can be described by using two gamma distributions (shown as the plots in blue color). The subplots show the similarity between the empirical cumulative probabilities and the cumulative probabilities of corresponding fitted gamma distributions. The one-sample Kolmogorov-Smirnov test is applied to verify the fitness of the gamma distribution.

## Discussion

Given the critical roles of spiking-timing patterns in synaptic plasticity and neural coding, it is important to understand the statistical patterns in spike-discharge dynamics in a brain-wide manner, as well as across multiple mammalian species. Although researchers have extensively modeled spike-timing patterns over decades of research, only several studies have compared spike regularity or irregularity over multiple brain subregions and/or animal species (Maimon and Assad, 2009; Mochizuki et al., 2016). One major caveat of any comparative experiment is that datasets were collected using rather different recording methods and with or without stringent spike-sorting criteria, leading to variations that may affect the outcome. To circumvent or minimize such technical variability, we have employed large-scale tetrode recording techniques and recorded from a total of 10 brain regions from mice and one region from hamsters using the same procedures and similar experimental protocols. These datasets were further compared with additional datasets from two brain regions of cats and monkeys that were collected by other laboratories. This systematic analysis of a wider range of brain regions, behavioral states, cell types and animal species has produced several important insights that are revealing to us.

First, the gamma distribution consistently outperforms Poisson distribution or log-normal distribution models across all datasets. Throughout the literature, spike patterns were often modeled with three different methods - namely, gamma distribution (Kuffler et al., 1957; Averbeck, 2009; Maimon and Assad, 2009; Mochizuki et al., 2016), log-normal distribution (Hromadka et al., 2008; Mizuseki and Buzsaki, 2013a) and Poisson-distribution process (Amarasingham et al., 2006; Beck et al., 2008), respectively. Because three statistical models have made different assumptions with underlying biological meaning that may or may not be true, it would be important to assess the statistical patterns over a wide range of cell types, regions, brain states and animal species, which have not been performed until now. We found that the neuronal spike-timing patterns have all conformed to gamma distributions.

In our analyses, we showed that Poisson process can only account for very small numbers of cells (when shape parameter *k*=1) (see Figure 13), a special case of the gamma-distribution. This is consistent with the notion that the Poisson model can either underestimate or overestimate the variabilities of neuronal spike patterns (Kara et al., 2000; DeWeese et al., 2003; Lindner, 2006; Heil et al., 2007; Kang et al., 2010; Berkes et al., 2011; Li et al., 2015; Moezzi et al., 2016). According to the independent spike hypothesis that the generation of each spike is independent of all the other spikes, the neuronal spike train would be completely described as a particular kind of random process called the Poisson process. If the Poisson process holds true for neuronal spike-activity patterns, ISI distribution would be statistically described by the exponential-distribution model. However, there are certain important features of neuronal firing that may render this simplistic assumption unsatisfactory or inaccurate for capturing spike-time patterns. In fact, burst-firing and the refractory periods after the generation of spikes can make neuronal spike patterns deviate away from a Poisson process (exponential-distribution model). We showed that the neuronal spike-timing patterns were best captured by gamma distributions.

Recently, log-normal distribution has been proposed to account for many variables of brain parameters (e.g., firing-rate distributions of neurons, the number of synaptic contacts between neurons, and the size of dendritic boutons), which had a positive-skewed, long-tailed distribution (Mizuseki and Buzsaki, 2013; Buzsaki and Mizuseki, 2014). Quantifications of cortical principal neurons have shown that the mean firing rates of individual neurons can span at least four orders of magnitude and that the firing-rate distribution of both stimulus-evoked and spontaneous activity in cortical neurons fit well with the log-normal distribution model. While our results support the notion that spike-timing patterns of single neurons followed a positive-skewed distribution, we found that the gamma-distribution model has consistently offered more precise fitting to the spike-timing patterns from all these brain regions, cell types, animal species and brain states.

Second, assessing from 13 different brain regions, we noted that there is the lack of a clear trend in increased regularity from primary sensory to higher cortical or motor cortical areas, as two previous studies had indicated (Maimon and Assad, 2009; Mochizuki et al., 2016). While our analysis of the parietal cortex dorsal area 5 (area 5d) in the monkey brain indeed found these motor-related neurons exhibited higher regularity, we showed that the mouse M1 cortex seemed to have similar irregular firing patterns in comparison to the mouse V1, auditory cortex, and RSC - as well as the prefrontal cortices, such as the PrL and ACC. The fact that we confirmed more regular firing in monkey 5d datasets from Cui laboratory argue against the idea that the discrepancy was simply due to different recording and analysis techniques. One possible explanation for this discrepancy between monkey parietal cortex area 5d and mouse M1 cortex is the difference in how the datasets were collected under different training conditions. In monkey, spike datasets were typically collected after monkeys were subjected to extensive behavioral training over many weeks and months, whereas our M1 datasets from mice were collected from naïve mice which were not subjected to any particular motor tasks. In fact, past studies have reported that cortical neurons, including M1 neurons, can generate spike trains with high irregularity (bursting) (Mountcastle et al., 1969; Bair and Koch, 1996; Buracas et al., 1998; Masse and Cook, 2008). Thus, trial conditions (such as oscillatory stimulus presentation or rhythmic motor actions) and extensive training of animals (over weeks and months) on motor tasks may influence spiking dynamics via experience-dependent plasticity and learning experience.

Among cells exhibiting higher firing regularity, we noted that DA neurons in the VTA and excitatory principal cells from the somatosensory cortex of the mouse brain belonged to this list. Tonic firing of DA neurons with their unique synaptic and cellular machinery can contribute to this unique pattern. We and other laboratories have reported different subtypes of DA neurons (Wang and Tsien, 2011), it will be interesting to further analyze how subtype DA neurons differ in their spike-timing patterns in future experiments.

We found that principal neurons in the mouse and cat V1 cortex – such as principal excitatory cells from the mouse retrosplenial cortex, secondary auditory cortex, anterior cingulate cortex, basolateral amygdala and hippocampal CA1 pyramidal cells – as well as inhibitory projection cells of striatum and fast-spike putative local interneurons, all tend to exhibit greater irregularity, a property likely due to different burst-firing dynamics of these neurons. It is noteworthy that current classification of cell types were based on waveforms and/or firing rates. More sophisticated classification will be necessary for future analyses. It is known that even within CA1 pyramidal cells, they may contain distinct subtype cells with a different burst index (Mizuseki et al., 2011; Li et al., 2015), a feature that can affect spike-time irregularity.

Third, by taking advantage of recorded spike datasets collected from the same animals under different cognitive states, we have examined the question of how different brain states influence the overall statistical patterns of spike-timing dynamics. Accordingly, we have verified that the spike-timing patterns conformed to gamma distribution under the quiet-awake and sleep states. One may argue that the spike-timing patterns during these brain states could be represented or dominated by “noise”. Therefore, it would be important to exclude such a possibility by using spike datasets from animals during more active states. By performing fearful stimulation experiments, we showed that gamma-distribution spike patterns not only continued to remain robust, but also the gamma-distribution model provided even more accurate-fitting results in comparison to awake resting or sleep states.

It should be noted that despite neurons’ mean firing rates varied between distinct natural states (i.e., sleep vs. awake resting period) (shown by the scale parameter θ in the right subpanels of Figures 5A-D), the overall shapes of the neuronal spike patterns are preserved across different states (shown by the scale parameter *k* in the left subpanels of Figures 5A-D). This supports the notion that while distinct brain activity states or inputs may boost or lessen the mean firing rates, the statistical characteristics of spike-timing patterns are able to remain robust and likely reflect preconfigured, skewed synaptic connections and/or intrinsic electrical/chemical attributes of a neuron. Similarly, the datasets from the anaesthetized cat V1 under two different visual stimulation conditions also showed the conserved gamma-distribution patterns. This conserved characteristic may be useful in term of offering a potential way of estimating a neuron’s spike pattern among distinct states. For example, instead of recording a single neuron’s spike train for a long period to get the statistical distribution of its spike pattern in the awake resting state, one may predict it by using the shape parameter *k* calculated under the sleep state and/or a mean firing rate within a shorter period in the stimulated state, or vice versa.

Fourth, given the widely conserved gamma-distribution spike-timing patterns across cell types, brain regions and naturally-occurring cognitive states, one may ask how important this pattern would be. In other words, does the disruption of this conserved gamma distribution pattern alter brain state and/or cognition? To address this question, we used ketamine-induced general anesthesia as a way to examine the relationship between gamma-distribution patterns and cognitions. The main action of ketamine is the blockade of the NMDA receptors, which is traditionally believed to reduce activation in the thalamo-cortical structures via blocking the inputs of afferent stimuli and information integration to the conscious mind (Alkire et al., 2008; Sinner and Graf, 2008). Interestingly, we found that ketamine turned a singular gamma distribution of spike-time patterns in the BLA, CA1 and RSC cells into two distinct subtypes of gamma distributions (see gamma distributions 1 and 2 in Figure 14). Most notably, subtype-2 gamma distribution showed highly regular firing patterns, which seem to be associated with the emergence of the delta band oscillation in simultaneously recorded LFP. Therefore, ketamine-induced unconsciousness is different from natural unconscious (sleep) states in term of dynamic spike-timing patterns.

In conclusion, we systematically investigated the statistical patterns of spike-timing irregularity in 13 different cortical and subcortical regions of mouse, hamster, cat and monkey brains. By comparing the effectiveness of different models, we show that spike-timing patterns of various projection neurons - including cortical excitatory principal cells, hippocampal pyramidal cells, inhibitory striatal medium spiny neurons and dopaminergic neurons, as well as fast-spiking interneurons – all invariantly conform to the gamma-distribution model. While higher regularity in spike-timing patterns is observed in a few cases, such as mouse DA neurons and monkey motor cortical neurons, there is no clear tendency in increased firing regularity from the sensory and subcortical neurons to prefrontal or motor cortices. Moreover, gamma shapes of spike-timing patterns remain robust over various natural brain states - such as sleep, awake resting periods, as well as during fearful experiences. However, ketamine-induced general anesthesia led to the breakdown of spike patterns from a singular gamma distribution into two distinct subtypes of gamma distributions. Taken together, the above results suggest that gamma-distribution patterns of spike timing reflect not only a fundamental property (Song et al., 2000; Lisman and Spruston, 2005) conserved across different neurons, regions and animal species, but also an operation crucial for supporting naturally-occurring cognitive states. Such gamma-distribution-based, spike-timing patterns may also have important implications for investigating real-time neural coding (Buzsaki, 2010; Gupta et al., 2010; Xie et al., 2016a) and perhaps better modeling of neuromorphic computing.

## Materials and Methods

### Ethics Statement

All animal work described in the study was carried out in accordance with the guidelines laid down by the National Institutes of Health in the United States, regarding the care and use of animals for experimental procedures, and was approved by the Institutional Animal Care and Use Committee of Augusta University (Approval AUP number: BR07-11-001).

### Construction of tetrode headstages and animal surgery of the datasets of mouse BLA, CA1, ACC, RSC, STR, V1, 2^nd^ AuV, M1, and somatosensory cortex, and hamster PrL

Tetrodes and headstages were constructed using the procedures as we have previously described (Lin et al., 2006; Xie et al., 2016a; Xie et al., 2016b). To construct tetrodes, a folded piece consisting of four wires (90% platinum, 10% iridium, 13 μm, California Fine Wire Company, Grover Beach, CA, USA) was twisted together using a manual turning device and soldered with a low-intensity heat source (variable temperature heat gun 8977020, Milwaukee, Brookfield, WI, USA) for six seconds. The impedances of the tetrodes were measured with an electrode impedance tester (Model IMP-1, Bak Electronics, Umatilla, FL, USA) to detect any faulty connections, and our tetrodes were typically between 0.7 MΩ and 1 MΩ. The insulation was removed by moving the tips of the free ends of the tetrodes over an open flame for approximately three seconds. The tetrodes were then placed into appropriate polyimide tubes. The recording ends of the tetrodes were cut differentially (Vannas spring scissors −3 mm cutting edge, Fine Science Tools, Foster City, CA, USA) according to the different depths of the recording sites. This ensures that only tetrodes, but not the surrounding polyimide tubes, were inserted into the brain tissue, thereby minimizing the tissue damage.

We employed adjustable 128-channel tetrode microdrives to target the basolateral amygdala (BLA; n = 8 WT mice), hippocampal CA1 (n = 9 WT mice), anterior cingulate cortex (ACC; n = 7 WT mice), the retrosplenial cortex (RSC; n = 20 WT mice), dorsal striatum (STR; n = 7 WT mice), primary visual cortex (V1; n = 14 WT mice), 2^nd^ auditory cortex (2^nd^ AuV; n = 9 WT mice), somatosensory cortex (n = 11 WT mice), and hamster prelimbic region (PrL; n = 13 WT Golden Syrian hamsters) bilaterally with 64 channels per hemisphere (Lin et al., 2006). Stereotaxic coordinates were as follows: for BLA, 1.7 mm posterior to bregma, 3.5 mm lateral, −4.0 mm ventral to the brain surface; for ACC, +0.5 mm AP, 0.5 mm ML, −1.75 mm DV; or CA1, 2.0 mm lateral to the bregma and 2.3 posteriors to the bregma; for RSC, −2.5 mm AP, 0.5 mm ML, −0.8 mm DV; for STR, +0.62 mm AP, 1.35 mm ML, −2.0 mm DV; for V1, −3.08 mm AP, 2.5 mm ML, −0.5 mm DV; for 2^nd^ AuV, −1.94 mm AP, 4.75 mm ML; for somatosensory cortex, −1.1 mm AP, 1.5 mm ML; and for recording in the prelimbic cortex (PrL) of the Golden Syrian hamster, the stereotaxic coordinate was +3.50 mm AP, ± 0.7 mm ML, −4.0 mm DV (Paxinos and Franklin, 2004).

Male wild-type mice (6–8 months old) or adult male hamsters (3–4 months old) were moved from home cages housed in the LAS facility to the holding area next to the chronic recording rooms in the laboratory and stayed in a large plastic bucket (20 inches in diameter and 16 inches in height – per mouse, Walmart) with access to water and food for a week prior to surgery. During this period, the animals were also handled daily to minimize the potential stress from human interaction. On the day of the surgery, the animal was given an intraperitoneal injection of 60 mg/kg Ketamine (Bedford Laboratories, Bedford, OH, USA) and 4 mg/kg Domitor (Pfizer, New York, NY, USA) prior to the surgery. The head of the animal was secured in a stereotaxic apparatus, and an ocular lubricant was used to cover the eyes. The hair above the surgery sites was removed, and Betadine solution was applied to the surface of the scalp. An incision was then made along the midline of the skull. Hydrogen peroxide (3% solution, Fisher Scientific) was placed onto the surface of the skull so that bregma could be visualized. The correct positions for implantation were then measured and marked. For fixing the microdrive headstage, four holes for screws (B002SG89S4, Amazon, Seattle, WA, USA) were drilled on the opposing side of the skull and, subsequently, the screws were placed in these holes with reference wires being secured to two of the head screws. Craniotomies for the tetrode arrays were then drilled, and the dura mater was carefully removed. After the electrodes were inserted and tetrodes were secured to the fiberglass base, the reference wires from the connector-pin arrays were soldered such that there would be a continuous circuit between the ground wires from the head screws and those from the connector-pin arrays. Finally, the connector-pin arrays were coated with epoxy. Aluminum foil was used to surround the entire headstage to aid in the protection and to reduce noise during recordings. The animals were then awoken with an injection of 2.5 mg/kg Antisedan. The animals were allowed to recover post-surgery for at least 3–5 days before recording began. Then, the electrode bundles targeting the BLA, STR and hippocampal CA1 region were slowly advanced over several days in small daily increments. For the cortical sites, tetrodes were advanced usually only once or twice in a small increment. At the end of the experiments, the mice were anesthetized, and a small amount of current was applied to the recording electrodes in order to mark the positions of the stereotrode bundles. The actual electrode positions were confirmed by histological Nissl staining using 1% cresyl echt violet. In some experiments, for facilitating the identification of electrode array position, the electrode tips were dipped in fluorescent Neuro-Dil (Neuro-Dil, #60016, Red oily solid color, from Biotium, Inc.), which then reveal the electrode track.

### Experimental Design

We recorded mouse (B6BCA/J) STR, V1, 2^nd^ AuV, M1, somatosensory cortex, and hamster PrL when animals were in the quiet-awake state. The datasets of the mouse (B6BCA/J) ACC, RSC, CA1 and BLA were recorded under two distinct states: the quiet-awake state and sleep state.

### *In vivo* recording of the datasets of the mouse BLA, CA1, ACC, RSC, STR, V1, 2^nd^ AuV, M1, somatosensory cortex, and hamster PrL

After surgery, the animals were handled for another 5–10 days while electrodes were advanced to the recording sites for obtaining the maximum neural units.

For the datasets of the mouse STR, 2^nd^ AuV, M1, somatosensory cortex, and hamster PrL, recordings were carried out while the animals were in their home cages. Quiet-awake episodes were manually assigned when the mice were awake and immobile. Neuronal spike data during the awake state was recorded for at least 15 minutes for each animal.

For the datasets of the mouse ACC, RSC, CA1 and BLA, we recorded the neuronal spike activity in both the quiet-awake state and the sleep state. Recordings were first carried out while the animals were in their home cages. To identify the sleep episodes, local field potentials were first band-pass filtered in theta band (4–12 Hz) and delta band (1– 4 Hz); then the ratio of the power in theta band to that of delta band was calculated. Two criteria were applied to extract from the sleep state: (1) Duration of an epoch was longer than five seconds, and (2) the ratio of the power during an epoch was greater than mean 5SD. For each mouse, the awake and sleep states were recorded for at least 15 minutes.

For the dataset of mouse V1, we set up an experimental configuration modified from the design of a previous study (Niell and Stryker, 2010). Briefly, a treadmill (five inches in diameter) was levitated by a stream of pressured air, the tangential force of a mouse running on the treadmill was calculated to be comparable to the force needed for free running. There was a lick port in front of the mouse’s mouth, dropping sugar water every minute. A camera with a telescope was set at the right side of the mouse eye (8cm distance) for monitoring eye movement. The mouse has implanted electrode arrays in the primary visual cortex of the left hemisphere for chronic recording a week before a head-fixed visual stimulation experiment. For habituation, the mouse was fixed on the top of a treadmill two hours every day, by mounting the headstage on a metal holder. An optical mouse was placed underneath the treadmill to measure the animal’s movements. During the recording, the mouse was to either sit still or run on the treadmill. We found that the eye movement and animal’s movement state were highly synchronized; the eyeball was almost fixed when the mouse was in a still state.

Visual stimuli were generated in MATLAB and displayed on a Dell 24-inch monitor placed 27 cm in front of the right eye. Before any recordings, we calibrated the location of the monitor. Specifically, black and white dots with one - four degrees were randomly presented on the monitor with a 10 ms duration. The visual responses of the primary visual cortex were amplified, acquired and streamed to the computer by the Plexon electrophysiology recording system. The time stamps of the visual stimulations were also sent to the Plexon simultaneously. The receptive field mapping software was written in MATLAB, which could tag the receptive field of the recorded neurons on the monitor, based on the e-Phys data. Then we adjusted the location of the monitor to make sure the neuron group receptive field would be located in the center of the monitor.

Two different visual stimuli were employed in the formal recording: 1. Full-length drifting bars [combinations of eight orientations (π/8, π/4, π3/8, π/2, π5/8, π3/4, π7/8 and π), four spatial frequencies (0.25, 0.5, 1 and 2 Hz), four temporal frequencies (0.25, 0.5, 1 and 2 Hz) and two directions (drifting either left or right)]; and 2. Two-dimensional stimuli [we recorded 624 gray-scale images of 78 objects (42 toy animals and 36 toy cars), each object was recorded from eight different viewing directions in a 45º interval]. For each recording section, one of these three visual stimuli were delivered. At least 30 minutes of neuronal spike data were recorded during each recording section. Basically, the mice stayed on the treadmill without struggling for more than two hours, then they were freed to the home cages.

### Cell-type classification in mouse BLA, ACC, RSC, STR, CA1 and hamster PrL

For the datasets recorded from the mouse BLA, ACC, RSC, STR, CA1 and hamster PrL, well-isolated units were classified as either putative excitatory principal cells or inhibitory interneurons based on three characteristic features of their spike activities - namely, trough-to-peak width, half-width after trough, and the mean firing rates. The k-means method was employed to achieve automated cell-type clustering. In general, putative principal cells fire at lower rates and have broader waveforms, whereas interneurons have higher rates and relatively narrower waveforms.

### Animal surgery and *in vivo* recording of mouse VTA dataset

The details have been previously described (Wang and Tsien, 2011). A 32-channel (a bundle of eight tetrodes) electrode array was constructed. One week before surgery, mice (3–6 months old) were removed from the standard cages and housed in customized home cages (40×20×25 cm). On the day of surgery, the mice were anesthetized with Ketamine/Xylazine (80/12 mg/kg, i.p.); the electrode array was then implanted toward the VTA in the right hemisphere (3.4 mm posterior to bregma, 0.5 mm lateral and 3.8–4.0 mm ventral to the brain surface) and secured with dental cement.

Two or three days after surgery, the electrodes were screened daily for neural activity. If no dopamine neurons were detected, the electrode array was advanced 40∼100 µm daily, until we could record from a putative dopamine neuron. In brief, spikes (filtered at 250–8000 Hz; digitized at 40 kHz) were recorded during the entire experimental process using the Plexon multichannel acquisition processor system (Plexon Inc.). Mice behaviors were simultaneously recorded using the Plexon CinePlex tracking system. Recorded spikes were isolated using Plexon Offline Sorter software. Dopamine neurons were distinguished from other neurons in the region by the characteristics of their extracellularly recorded impulses - including long, multiphasic waveforms and low basal firing rates.

### Fearful-event experiment

Mice were subjected to three fearful episodic events, earthquakes, foot shocks, and free-fall drops. For the earthquake-like shake, the mouse was placed in a small chamber (4″ × 4″ × 6″H circular chamber) fixed on top of a vortex mixer and shaken at 300 rpm for 400ms six times with 1~3-minute time intervals between each shake. For fearful foot shock, the foot-shock chamber was a square chamber (10″ × 10″ × 15″H) with a 24-bar shock grid floor. The mouse was placed into the shock chamber for three minutes and received the foot-shock stimulus (a continuous 300-ms foot shock at 0.75 mA) for a total of six times with inter-trial time interval between 1~3 minutes. For free-fall in the elevator, the animal was placed inside a small box (3″ × 3″ × 5″H) and dropped from a 13-cm height (a cushion which made from a crumbled tablecloth was used to dampen the fall and to stop the bouncing effect). After 1~2 minutes, the elevator was raised gently back to the 13-cm height and dropped again after 1~2 minutes (this process was also repeated six times). These episodic stimuli are fearful as evidenced from physiological indications including a rapid increase in heart rates as well as reduced heart rate variability (Liu et al., 2013; Liu et al., 2014). To maintain the consistency of stimulation timing (minimizing the possible prediction of upcoming stimuli), the stimuli were triggered by a computer and delivered at randomized intervals within 1-3 minutes. After the completion of all fearful event sessions, the mouse was placed back into the home cage.

### Dataset recorded from the quiet-awake state and anesthesia

Three datasets were recorded from the BLA, hippocampal CA1, and RSC under two distinct states - namely the quiet-awake state and anesthesia. Recordings were first carried out when animals were awake and immobile in their home cages. Minimum 40-minute neural activities were recorded in the quiet-awake state. To produce ketamine-induced anesthesia, the animals were injected with a 60mg/kg Ketamine and 0.5 mg/kg Domitor cocktail mixture via *i.p.*; the animals lost the righting reflex in a few minutes. Neural spike activities were recorded for 50 minutes under the anesthetized state. Forty-minute neural spike data recorded from the fully anesthetized state starting from 10 minutes after the Ketamine/Domitor injections were selected for the present analysis. The dataset for neuronal variability analyses of awake state vs. anesthesia (Figure 14) contained 92 BLA putative principal cells from two mice, 59 CA1 putative principal cells from three mice, and 59 RSC putative principal cells from three mice.

### Monkey area 5d dataset

The dataset was previously described (Li and Cui, 2013). Data were recorded from two male rhesus monkeys (Macaca mulatta, 7-10 kg). The animals were trained to perform visually guided single- and double-arm reaching tasks. All procedures were in accordance with NIH guidelines and were approved by the Institutional Animal Care and Use Committee of Augusta University. Recorded spikes were re-sorted using Plexon Offline Sorter software for obtaining the spike trains of single neurons; multiple spike-sorting parameters (e.g., principal component analysis, energy analysis) were used for the best isolation of single-unit spike trains. Because of the absence of the spike waveform information, we were not able to separate different neuron types (excitatory vs. inhibitory cells, etc.). However, we observed that the previously-held conclusion that neuronal spike activities can be best described by the gamma-distribution model still holds true in the monkey area 5d region.

### Cat V1 Datasets

The cat V1 datasets (1-D and 2-D) were downloaded from the Collaborative Research in Computational Neuroscience (CRCNS) website (data from Yang Dan Lab at UC-Berkeley, download link: http://crcns.org/data-sets/vc/pvc-2). Experimental procedures have been previously described (Touryan et al., 2002; Felsen et al., 2005; Touryan et al., 2005). The data were obtained with extracellular recordings from the primary visual cortex of anesthetized adult cats. Single-unit recordings were made in area 17 of adult cats (2–6.5 kg) using tungsten electrodes (A-M Systems, Carlsborg, WA). Animals were initially anesthetized with isoflurane (3%, with O_2_) followed by sodium pentothal (10 mg/kg, i.v., supplemented as needed). During recording, anesthesia was maintained with sodium pentothal (3 mg · kg^−1^ · hr^−1^, i.v.), and paralysis was maintained with pancuronium bromide (0.1–0.2 mg · kg^−1^ · hr^−1^, i.v.). Because of the absence of the spike waveform information, we were not able to separate different neuron types (excitatory vs. inhibitory cells, etc.). However, we observed that the previously-held conclusion that neuronal spike activities can be best described by the gamma-distribution model still holds true in the cat V1 region (both 1-D and 2-D stimuli).

### Data processing and spike sorting

Neuronal activities from mouse experiments were recorded by the MAP system (multi-channel acquisition processor system, Plexon Inc., Dallas, TX) in the manner as previously described (Kuang et al., 2010). Extracellular action potentials and local field potentials data were recorded simultaneously and digitized at 40 kHz and 1 kHz, respectively. The artifact waveforms were removed, and the spike waveform minima were aligned using the Offline Sorter 2.0 software (Plexon Inc., Dallas, TX), which resulted in more tightly clustered waveforms in principal component space. Spike sortings were done with the MCLUST 3.3 program with an auto-clustering method (KlustaKwik 1.5). Only units with clear boundaries and less than 0.5% of spike intervals within a 1 ms refractory period were selected. The stability of the *in vivo* recordings was judged by waveforms at the beginning, during and after the experiments. Well-separated neurons were assessed by “Isolation Distance” (Schmitzer-Torbert et al., 2005). Neurons whose “Isolation Distance” >15 were selected for the present analysis as previously described (Lin et al., 2006; Xie et al., 2016a; Xie et al., 2016b).

### Statistical Analysis

As shown in Figure 5, a *t*-test was used to assess whether *k* (or *θ*) was linearly correlated between the quiet-awake state and the sleep state. In Figure 11, one-way ANOVA analysis and Tukey post-hoc tests were conducted for the comparisons of multiple means of *k* across different brain regions. Three asterisks denoted the p-value is less than 0.001. Data were represented as mean ± SEM.

### Statistical properties of neural spike patterns

A two-step analysis was employed to examine the statistical properties of neural spike patterns.

**Step 1**: Positive-skewed vs. negative-skewed distributions. The distribution of ISI was characterized by two well-defined statistics - namely, nonparametric-skew (*S*) and skewness (*γ*), defined as follows:

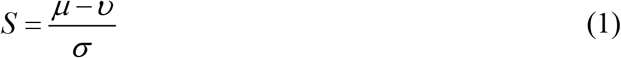

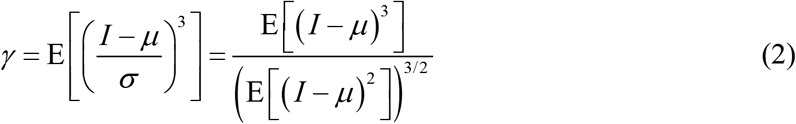

where *μ* is the mean, *υ* is the median, *σ* is the standard deviation, E is the expectation operator, and *I* denotes the ISIs of a neuronal spike train.

In probability theory and statistics, nonparametric-skew (*S*) and skewness [*y* ∊ (-∞, +∞)] are measurements of the skewness (or long-tailedness) of a random variable’s distribution - that is, the distribution’s tendency to lean to one side or the other of the mean (Figure 1B). A positive-skewed distribution (red curve in Figure 1B) has *S* > 0 and *γ* > 0, a negative-skewed distribution (blue curve in Figure 1B) has *S* < 0 and *γ* < 0, while a symmetric distribution (gray dotted curves in Figure 1B) has *S* = 0 and *γ* = 0.

**Step 2**: Gamma distribution vs. log-normal distribution. Let *I*_1_,*I*_2_,…*I*_n_ denote the ISIs of a neuron’s spike train, the probability density function for a gamma distribution of *I* is defined by a shape parameter *k* > 0 and a scale parameter *θ* > 0:

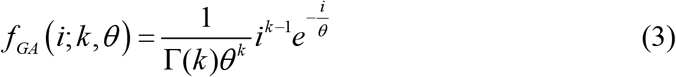

where Γ(*k*) is the gamma function.

The probability density function of a log-normal distribution of *I* is defined by a location parameter *μ* ∊ (−∞, +∞) and a scale parameter *σ* > 0:

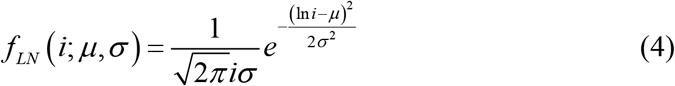

In the present analyses, both gamma distribution and log-normal distribution were estimated using the maximum likelihood estimates (MLE) method, which selected values of parameters that produced corresponding distributions for the histograms of ISI with minimal errors. The parameters of both gamma distribution and log-normal distribution were directly estimated by using the raw ISIs of neuronal spike activities, which ensured that the analyses of ISI distributions were independent of the bin sizes used.

Goodness-of-fit analyses were then conducted to quantitatively discriminate between the gamma-distribution and log-normal distribution models. For convenience, we termed a gamma distribution and a log-normal distribution as *GA*(*k,θ*) and *LN*(*μ,σ*), respectively. Thus, the likelihood functions of data that follow *GA*(*k,θ*) and *LN*(*μ,σ*) can be denoted as:

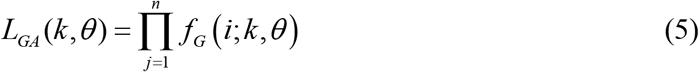

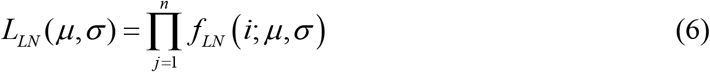

Therefore, the RML between a gamma distribution and a log-normal distribution is defined as:

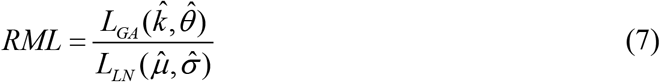

where 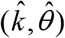 and 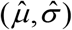 are the MLEs of (*k,θ*) and (*μ,σ*) for *I*. The natural logarithm of RML can be written as:

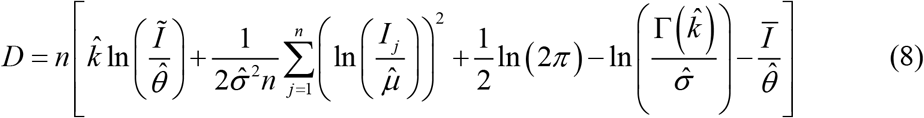

where 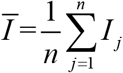 and 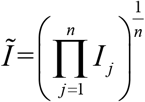 are the arithmetic and geometric means of *I*, respectively.

The natural logarithm of RML, *D*, measured the goodness-of-fit between the gamma-distribution model and the log-normal distribution model of ISI. That is, the gamma-distribution model precedes the log-normal distribution model if *D* > 0; otherwise, choose the log-normal distribution as the preferred model of ISI. The larger the absolute value of *D*, the more accurate-fitting the result of the chosen model over the other model.

### Local field potential spectral analysis

To generate the power spectra of LFP in Figure 14, the local field potential power density was calculated in a range of 0.025–20.0 Hz with 0.025 Hz intervals for the recording dataset of hippocampal CA1, BLA and RSC regions. The fast Fourier transform (with Hann windowing function) was applied to the EEG signal, the resulting frequency resolution was 0.025 Hz, and the frequency bins less than 1 Hz were discarded due to the sensitivity of these bins to noise.

## Author Contributions

J.Z.T. and M.L. conceived and designed the project. J.Z.T. designed the experiments with M.L., K.X., H.K., G.E.F. and J.L. The research was performed as follows: K.X. recorded from the ACC and PrL datasets; G.E.F. from the RSC; Jun Liu from the BLA; D.W. from the striatum, H.K. for the CA1, M.L., F.Z., Z.S., L.C., Y.M., and H.K. for data analyses together with J.Z.T. J.Z.T. and M.L. wrote the paper with input from all others.

## Acknowledgements

This work is supported by an NIH grant (R01NS079774) and GRA equipment grant to J.Z.T., Shanghai Youth Science and Technology Sail Project (16YF1415200) to Z.S. and M.L., the Brain Decoding Center grant Yunnan Science Commission (2014DG002) to F.Z. and J.Z.T. We thank Prof. Yang Dan at UC Berkeley for sharing the cat dataset, which was downloaded from the CRCNS website. We thank Sandra E. Jackson for proofreading the manuscript.

